# Topology-dependent DNA binding

**DOI:** 10.1101/2023.06.30.547266

**Authors:** Pauline J. Kolbeck, Miloš Tišma, Brian T. Analikwu, Willem Vanderlinden, Cees Dekker, Jan Lipfert

## Abstract

DNA stores our genetic information and is ubiquitous in biological and biotechnological applications, where it interacts with binding partners ranging from small molecules to large macromolecular complexes. Binding is modulated by mechanical strains in the molecule and, in turn, can change the local DNA structure. Frequently, DNA occurs in closed topological forms where topology and supercoiling add a global constraint to the interplay of binding-induced deformations and strain-modulated binding. Here, we present a quantitative model of how the global constraints introduced by DNA topology modulate binding and create a complex interplay between topology and affinity. We focus on fluorescent intercalators, which unwind DNA and enable direct quantification via fluorescence detection. Using bulk measurements, we show that DNA supercoiling can increase or decrease intercalation relative to an open topology depending on ligand concentration and the initial topology. Our model quantitatively accounts for observations obtained using psoralen for UV-induced DNA crosslinking, which is frequently used to quantify supercoiling *in vivo*. Finally, we observe topology-dependent binding in a single-molecule assay, which provides direct access to binding kinetics and DNA supercoil dynamics. Our results have broad implications for the detection and quantification of DNA and for the modulation of DNA binding in cellular contexts.

## INTRODUCTION

DNA is the carrier of genetic information in all cellular life. *In vivo*, double-stranded DNA is often present in circular and, therefore, topological closed form. In particular, bacterial chromosomes and plasmids are circular DNA molecules, whose topology is tightly regulated *in vivo* (1–4). In eukaryotes, DNA topology and supercoiling similarly play important roles in the context of a chromatinized genome, e.g., in the compaction and regulation of genetic information (5–9).

Both in its biological role and in many biotechnological applications, DNA interacts with a broad range of ligands, i.e., binding partners that range from small molecules to large proteins complexes. In particular, the detection and quantification of DNA often rely on staining with fluorescent small molecules that frequently bind in an intercalative binding mode (10–15). Ligand binding to DNA can, in general, locally alter the DNA structure and introduce strains away from the equilibrium B-form DNA conformation (16–20). In turn, stretching forces and torsional strains have been shown to systematically affect binding equilibria (21–25). Having a defined DNA topology, e.g., in a plasmid or other topological domains, imposes a global constraint on the interplay between strain-dependent binding and binding-induced conformational changes. However, it is not well understood how this interplay affects binding in a typical experimental setting and how to quantitatively model binding equilibria under a global topological constraint.

Here we investigate DNA-ligand binding under a global topological constraint. We first develop a model of ligand binding to topologically closed DNA, which takes into account the conformational changes induced by the ligand, the influence of strains on binding equilibria, a physical model of plasmid mechanics, and the global constraint introduced by having a defined linking number due to the defined topology. We then experimentally probe interactions between small molecule intercalators and DNA of different, defined topologies. We focus on commonly used intercalators that are well-characterized by previous studies: Ethidium bromide (EtBr), SYBR Gold, and SYTOX Orange (Supplementary Figure S1). EtBr is a very widely used stain for DNA visualization in gels and other applications (23–31). SYBR Gold is a more recently developed DNA stain, which has very high quantum efficiency and brightness (15,32–34). SYTOX Orange is frequently used to stain and supercoil DNA in single-molecule experiments (34–38). Finally, we extend our analysis to the intercalator 4,5’,8-trimethylpsoralen (TMP; also known as trioxsalen) that is used as a photo-crosslinking agent, both for phototherapy (39) and to detect supercoiling and chromatin structure *in vivo* (40–44).

We first perform experiments in bulk using topologically constrained plasmid DNA, i.e., circular DNA with both strands fully intact (referred to as the supercoiled species, “sc”; Figure 1A) and, for comparison, topologically open DNA that has been either nicked (open circular, “oc”; Figure 1B) or linearized (“lin”; Figure 1C). Additionally, we use fluorescence detection to quantify the amount of binding, and demonstrate that intercalative binding to DNA is topology dependent, in quantitative agreement with our model. We then apply our model to a single-molecule DNA assay (35,38), where DNA is supercoiled *in situ* by intercalation. We monitor the fluorescence change upon nicking of the DNA molecule, which induces an abrupt transition from a closed to an open topology, and we find excellent agreement with our model. The single-molecule assay enables us to observe the re-adjustment of the binding equilibrium upon change in topology in real time, which enables us to probe the dynamics of the process.

**Figure 1.**
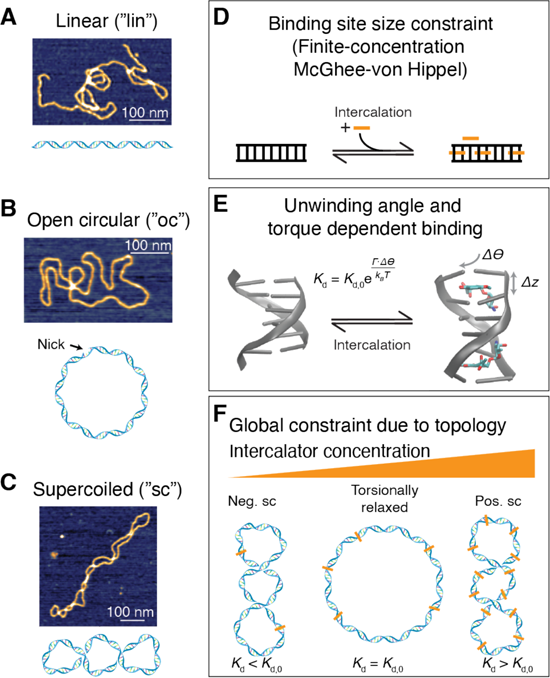
Overview of different topological conformations of plasmid DNA and outline of the binding model under global constraint. A) Schematic and an AFM height image of linear DNA. B) Schematic and an AFM height image of open circular DNA, i.e., of plasmid DNA that is nicked at a single site. C) Schematic and an AFM height image of a negatively supercoiled DNA plasmid. AFM images in panels A-C are of pBR322 plasmid DNA (4,361 bp; see Methods). D) Schematic of the McGhee-von Hippel binding model for ligand binding to DNA, whereby the binding site size modulates the binding equilibrium. E) Illustration of how intercalation into DNA lengthens and underwinds the B-form helix. Left: B-form DNA, rendered from PDB entry 4C64 (91). Right: DNA in the presence of an intercalator, here daunomycin rendered from PDB entry 1D11 (1). F) Schematic of how the global constraint due to topology leads to increased binding as long as the DNA is negatively supercoiled (left), but will decrease binding when the DNA is positively supercoiled (right), compared to the torsionally relaxed form (center). Intercalation, in turn, locally underwinds the DNA and, therefore, leads to an increase in *Wr* with increasing intercalation.

Our findings have direct practical applications since gel-based assays for the discrimination and detection of topoisomers are widely used to study the properties of circular DNA and of various enzymes that alter DNA topology, including topoisomerases, gyrase, reverse gyrase, and recombinases (45–47). An unbiased quantification of the different topoisomers using fluorescence staining, which is increasingly used to replace radiolabelling due to the hazards associated with handling, storing, and disposing of radioactive materials, must take into account the observed topology dependencies. Using our findings, we provide practical guidelines for unbiased detection of different topoisomers and discuss consequences of topology-dependent binding more broadly.

## MATRERIALS AND METHODS

### Plasmid DNA preparation

We used the plasmid pBR322 (NEB) as the DNA substrate for bulk measurements of intercalation. We prepared different topological states of the DNA by cutting (i.e. introducing a double-strand break) or nicking (i.e. a DNA single-strand break) the original supercoiled DNA to obtain linear and open circular DNA, respectively. Reactions were performed in NEBuffer 3.1 (NEB) using the enzymes EcoRV (NEB; incubation temperature 37 °C) to prepare linear DNA and Nt.BspQI (NEB; incubation temperature 50 °C) to create open circular DNA. The reactions were stopped after one hour by heat inactivation at 80 °C for 20 minutes. The products were purified with a PCR clean-up kit (Qiagen). The concentrations were determined using a nanodrop UV/vis photospectrometer (ThermoFisher Scientific). To every 100 μl reaction volume, we added 20 μl Gel Loading Dye Purple (6x) (NEB, catalogue number B7024S) prior to running the gel. For the plate reader and qPCR cycler experiments, no gel loading dye was added.

### DNA dilution series

For the DNA dilution series, the different topologies were combined to an equimolar mixture and diluted with TAE buffer (40 mM Tris, 20 mM acetic acid, and 1 mM EDTA, pH 8.6) and gel loading dye in form of a serial dilution to obtain 7 different DNA concentrations (Table S1).

For the experiments using relaxed DNA, the plasmid pBR322 was relaxed using Wheat Germ Topoisomerase I (Inspiralis). For a total volume of 100 μl, 16.2 ul assay buffer (50 mM Tris HCl, 1 mM EDTA, 1 mM DTT, 20% (v/v) glycerol, 50 mM NaCl, pH 7.9), 82.12 μl RNase-free water, 2.16 μl topoisomerase I, 0.52 μl pBR322 (c = 1000 ng/μl) were combined in a reaction tube and incubated for 1 h at 37 °C. For the DNA dilution series, only the linear and the relaxed topologies were combined to an equimolar mixture and diluted with Tris-acetate-EDTA (TAE) buffer and gel loading dye in form of a serial dilution to obtain 7 different DNA concentrations (Table S1).

### AFM imaging of DNA plasmids

AFM imaging of plasmid DNA was performed as described previously (48–50). In brief, for the AFM imaging, we deposited 20 μl of DNA at different topological states in TE buffer at a final concentration of 1 ng/μl on freshly cleaved poly-L-lysine (Sigma Aldrich, diluted to 0.01% in milliQ water; PLL)-coated muscovite mica. The sample was incubated 30 s before washing with 20 ml MilliQ water and drying with a gentle stream of filtered argon gas. After drying, the AFM images were recorded in tapping mode at room temperature using a Nanowizard Ultraspeed 2 (JPK, Berlin, Germany) AFM with silicon tips (FASTSCAN-A, drive frequency 1400 kHz, tip radius 5 nm, Bruker, Billerica, Massachusetts, USA). Images were scanned over different fields of view with a scanning speed of 5 Hz. The free amplitude was set to 10 nm. The amplitude setpoint was set to 80% of the free amplitude and adjusted to maintain a good image resolution. AFM image post-processing was performed in the software SPIP (v.6.4, Image Metrology, Hørsholm, Denmark) to flatten and line-wise correct the images (Supplementary Figure S2).

### Gel electrophoresis

For gel electrophoresis we used 1%-broad-range-agarose (Carl Roth) gels and TAE buffer. We used the 1 kb gene ruler (Thermo Scientific; 5 μl) as a size standard. The gels were run for 120 min at 75 V at 4 °C. Subsequently, the gel were removed from the gel box and placed for 20 minutes in 100 ml of 0.5 μM (1:100000 dilution of the stock), 5 μM (1:10000 dilution of the stock), or 50 μM (1:1000 dilution of the stock) EtBr in TAE buffer, respectively, for staining. Subsequently, the gel was de-stained in TAE buffer for 15 min. The gels were visualized using a Gel Doc XR+ system (Biorad). Since the dye concentration used in staining is reduced by the agarose gel matrix and the de-staining step, we used a staining correction factor of 0.1 as determined previously (15), which we use to correct all gel data.

### Gel electrophoresis image analysis

We saved the images from the Gel Doc system in scn format to allow for quantitative fluorescence intensity analysis. The software SPIP (v.6.4, Image Metrology, Hørsholm, Denmark) was used to remove spikes from the image (without changing the intensity of the bands) and to generate average intensity profiles along each lane of the gel. In a next step, we used Origin (OriginLab, Northampton, Massachusetts, USA) to flatten the background of the profiles and to convert the peaks into fractions of supercoiled, open-circular, linear, and/or relaxed DNA respectively, by calculating the area under the lane profiles (Supplemenary Figure S3).

### Bulk fluorescence experiments

For the SYBR Gold bulk fluorescence measurements, we used a well plate reader (Tecan Infinite M1000 PRO; well plates: corning black polystyrene 384 well microplate with a flat bottom, Sigma-Aldrich, catalogue number: CLS3821) and a qPCR cycler (CFX96 Touch Real-Time PCR Detection System, BioRad). In the well-plate reader, the DNA mix including various SYBR Gold concentrations was filled in the wells and the fluorescence was read out from the bottom of the wells. The excitation and emission bandwidths were set to 5 nm, the gain to 100, the flash frequency to 400 Hz, and the integration time to 20 s. We chose the excitation and emission wavelengths - according to the excitation and emission maxima for SYBR Gold provided by Invitrogen - to be 495 nm and 537 nm.

In the fluorescence bulk experiments using a qPCR cycler, the DNA was filled into low-profile PCR tubes (Bio Rad, Nucleic Acids Research, 2021 5, product ID: TLS-0851), closed with flat, optical, ultra-clear caps (Bio Rad, product ID: TCS-0803) since the fluorescence was read out from the top of the tubes (at 24 °C). As read-out channels, channels with absorption and emission wavelengths of 494 and 518 nm, respectively, were chosen because these were the closest match to those of SYBR Gold.

### Single-molecule fluorescence experiments

Supercoiled DNA were prepared and imaged at the single-molecule level essentially as described previously (35,38). Experiments were performed in custom-made flow cells built by connecting a surface-passivated glass slide and a glass coverslip using double-sided tape (35). The surface of the glass slides was prepared as previously described (51) with slight modifications. In brief, after extensive cleaning, the surface was silanized using APTES (10% v/v) and acetic acid (5% v/v) methanol solution. The surface was passivated with NHS-ester PEG (5,000 Da) and biotinylated NHS-ester PEG (5,000 Da) in relation 40:1. The biotinylated NHS-ester PEG allowed us to tether the DNA molecules to the surface via biotin-streptavidin interactions (Fig. 5A).

The DNA used for intercalation with SYTOX Orange was prepared as described in Ref. (52) with the exception of introducing multiple-biotin handles at the DNA ends to allow the torsional constrain on the DNA rotation. To introduce multiple biotins on the DNA handles we performed a PCR on pBluescript SK+ (Stratagene) with GoTaq G2 DNA polymerase (Promega, M7845), in the presence of biotin-16-dUTP (Jena Bioscience, NU-803-BIO16-L) and dTTP (Thermo Fisher Scientific, 10520651) in molar ratio of 1:4, respectively. The PCR was done using primers: accgagatagggttgagtg and cagggtcggaacaggagagc, resulting in a 1,238 bp DNA fragment that contained multiple biotins randomly incorporated due to the presence of biotin-16-dUTP modified nucleotides. The PCR products were cleaned up using a standard purification kit (Promega, A9282) and we digested both the biotin handle and large 42 kb DNA plasmid with SpeI-HF (New England Biolabs, R3133L) for 2 h at 37 °C. The reaction was stopped by subsequent heat-inactivation for 20 min at 80 °C. This resulted in linear 42 kbp DNA and ∼600-bp biotin handles. The digested products were mixed, using a 10:1 molar excess of the biotin handle to linear plasmid. We then added T4 DNA ligase (New England Biolabs, M0202L) in the presence of 1 mM ATP overnight at 16 °C and subsequently heat-inactivated for 20 min at 65 °C. The resulting coilable 42 kbp DNA construct was cleaned up by size exclusion chromatography on an ÄKTA pure system, with a homemade gel filtration column containing approximately 46 ml of Sephacryl S-1000 SF gel filtration media (Cytiva), run with TE + 150 mM NaCl buffer. The sample was run at 0.2 ml min^−1^, and we collected 0.5 ml fractions.

For immobilization of this 42 kbp biotinylated-DNA, we introduced 100 µl of ∼0.5 pM DNA molecules at a flow rate of 4.2 µl/min in imaging buffer with no oxygen scavenging system components (40 mM Tris-HCl, 2 mM Trolox, 2.5 mM MgCl_2_, 65 mM KCl). The buffer during the DNA immobilization step contained either a low SYTOX Orange concentration (25 nM; to allow introducing positive supercoiling in a later step, see below), or high concentration of SYTOX Orange (250 nM; to allow introducing negative supercoiling in a later step, see below) during initial immobilization.

Immediately after the introduction of DNA molecules into the flow cell, we further flowed 100 µl of the same buffer without the DNA at the same flow rate to ensure stretching and tethering of the other end of the DNA to the surface (which introduces the topological constraint) as well as to remove unbound DNA molecules. To introduce plectonemic supercoils into the bound DNA molecules that are torsionally constrained, we change the SYTOX Orange concentration by introducing imaging buffer with the same salt concentration but with varying SYTOX Orange concentration and an oxygen scavenging system for imaging (40 mM Tris-HCl, 2 mM Trolox, 2.5 mM MgCl_2_, 65 mM KCl, 2.5 mM protocatechuic acid (PCA), 50 nM protocatechuate-3,4-dioxygenase (PCD)) (53). For positive supercoiling we increased the concentration from 25 nM during immobilization to 250 nM SYTOX Orange, while for the negative supercoiling we reduced the concentration from the initial binding at 250 nM to a final concentration of 50 nM for imaging. In both cases we checked for a visible presence of supercoiling on the DNA by moving foci appearing along the DNA molecules (Fig. 5, Supplementary Movies S1 and S2). Thus, supercoiling was introduced by differential SYTOX Orange concentrations between the DNA-binding step (where torsional constraint is ensured via multiple biotin molecules) and the imaging step where SYTOX Orange concentration is increased or reduced in order to generate positive or negative supercoils, respectively.

For fluorescence imaging, we used a home-built objective-TIRF microscope. We employed continuous excitation with a 561 nm (15-20 mW) laser in Highly Inclined and Laminated Optical sheet (HiLo) microscopy mode, to image SYTOX Orange-stained DNA as well as to introduce DNA nicking. All images were acquired with an PrimeBSI sCMOS camera at an exposure time of 20-200 ms, depending on the experiment, with a 60x oil immersion, 1.49NA CFI APO TIRF (Nikon). For DNA visualization, and kymograph generation, we used a custom written python software published in Ref. (54).

### Numerical implementation of the binding model under topological constraint

To model the effects of the global constraint imposed by DNA topology on binding, we developed a model that takes into account changes in linking number due to intercalation and, conversely, torque-dependent binding (see the section “Model for ligand binding under topological constraint” in Results). The model comprises coupled Equations 1-6 that are solved iteratively using a custom routine written in Matlab (Mathworks). Input parameters are the temperature *T* and for the DNA the number of base pairs *N_bp_*, the torsional stiffness *C,* the initial linking difference *ΔLk_0_,* and the DNA concentration *c*_DNA_*. N_bp_, ΔLk_0_*, and *c*_DNA_ are typically known from how the DNA was prepared and we used a value *C* = 100 nm, unless otherwise noted (55–57). For the single-molecule assay, the DNA concentration is poorly defined, but low, much lower than the intercalator concentration used. In this case, we used concentrations in the range of 1-100 pM·bp, which are much lower than the intercalator concentration used, but high enough to ensure numerical stability of the calculation. We found that the calculated results are insensitive to the concentration used in this regime.

Input parameters for the intercalator are the binding site size *n*, the dissociation constant *K*_d_, the length increase per intercalator bound Δz, the change in DNA helical twist per intercalator bound *Δθ*, and the intercalator concentration *c*_total_. Values for *n*, *K*_d_, *Δz*, and *Δθ* used in this work are provided in Table 1. The outputs of the model are the number of intercalator molecules bound per DNA molecule, the torque in the DNA, and the total fluorescence intensity expected, which contains an overall scaling factor *α* defined in Equation 7. Our numerical implementation uses the numerical solutions to the McGhee-von Hippel described previously (15,58) and a successive over-relaxation type approach (59) to speed up convergence.

**Table 1.** Parameters of selected intercalators used in this work. ^†^We used the values from Ref. (34) in 100 mM NaCl, close to the ionic strength used in this work. The value for *Δθ* is an estimate based on data for SYBR Gold. ^#^Ref. (89) only determined the dissociation constant approximately. The binding site size and elongation per dye for TMP were assumed to assume the average values for intercalators indicated in the table.

| Quantity | Ethidium<br>bromide<br>(EtBr)<br>Ref. (24) | SYBR<br>Gold<br>Ref. (15) | SYTOX<br>Orange<br>Ref. <sup>†</sup> (34) | Trimethylpsoralen<br>(TMP)<br>Ref. <sup>#</sup> (89) & (90) |
| --- | --- | --- | --- | --- |
| Binding site size $n$ | 1.9 | 1.6 | 3.0 | 2 |
| Dissociation<br>constant $K_d$ (M) | $7.7 \cdot 10^{-6}$ | $0.2 \cdot 10^{-6}$ | $0.4 \cdot 10^{-6}$ | $10^{-4}$ |
| Elongation per dye<br>$\Delta z$ (nm) | 0.34 | 0.34 | 0.30 | 0.34 |
| Untwisting per dye<br>$\Delta\theta$ (degree) | 27 | 19.1 | 19.1 | 28 |

## RESULTS

### Model for ligand binding under topological constraint

Here we develop a model for ligand binding to plasmid (or otherwise topologically closed) DNA, where the topology imposes a global constraint (Figure 1D-F). Binding to plasmid DNA is different from a standard bimolecular binding equilibrium for several reasons that we take into account in our model. First, the linear structure of DNA imposes local constraints for ligand binding, if bound ligands occupy a binding size of *n* bases (Figure 1D). Binding of ligands with binding site size *n* can be modeled using the McGhee-von Hippel model (24,60) in cases where the DNA concentration is much lower than the ligand concentration, such that the free and total ligand concentrations are approximately equal. For bulk measurements, however, the concentration of DNA bases can be similar to or even larger than the ligand concentration and needs to be considered. Therefore, we use an extension of the McGhee-von Hippel model that explicitly takes into account both the ligand (C*_total_*) and DNA (C*_DNA_*) concentration that was derived in Ref. (15). The fractional binding γ is given by

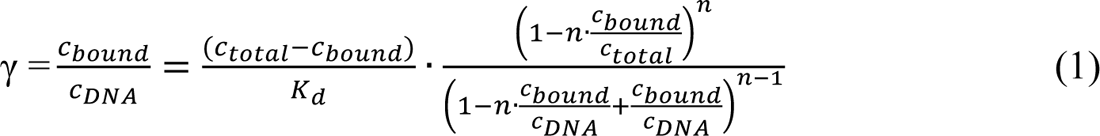

Here C*_bound_* is the bound ligand concentration, C*_free_* is the free ligand concentration, C*_total_* = C*_free_* + C*_bound_* the total ligand concentration, K*_d_* is the dissociation constant (in M), and *n* is the binding site size (in base pairs). Typical values of the binding site size for intercalators are *n* ≈ 2, corresponding to binding every other base pair.

Intercalation affects the local geometry of the DNA helix, by locally unwinding and lengthening the helix (11,12,23,24,34,61,62) (Figure 1E). Here we assume that each intercalation event locally lengthens the DNA by *Δz* and unwinds it by *Δθ*. Typical values for intercalators are in the range *Δz* ≈ 0.34 nm and *Δθ* ≈ 15°-30°. The fact that intercalation lengthens and unwinds the DNA helix suggests, by Le Chatelier’s principle, that applying a stretching force or unwinding torque, respectively will increase intercalative binding. Conversely, overwinding the helix will hinder intercalation. We assume an Arrhenius-like exponential dependence of the binding constant (23,30,34,63) on applied force *F* force and torque *Γ*:

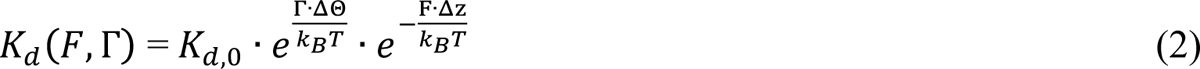

Here *K*_d,0_ is the dissociation constant for the relaxed molecule, i.e., in the absence of forces or torques, *k*_B_ Boltzmann’s constant and *T* the absolute temperature. For plasmids in free solution, the force is zero (or at least small, specifically *F* ≪ *k*_B_T / *Δz* ≈ 10 pN) and the second exponential factor in Equation 2 can be neglected. Values for *n*, *K*_d_, *Δz*, and *Δθ* for selected dyes are summarized in Table 1.

For linear or open circular DNA molecules, there is no torsional constraint, and the torque will be zero in equilibrium; consequently, binding will simply be determined by Equation 1. In contrast, for topologically closed plasmids, the topology imposes a global constraint. For a closed plasmid, the linking number *Lk* is a topological invariant and partitions into twist *Tw* and writhe *Wr* by White’s theorem (64):

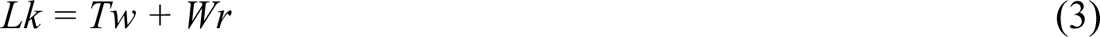

*Tw* is a measure for the local winding of the helix and directly related to the torsional strain in the molecule. Conversely, *Wr* corresponding the crossings of the double helical axis and in a plasmid is related to the number of plectonemic supercoils. We express the linking number balance relative to the torsionally relaxed double-stranded DNA, for which we define *ΔLk* = 0, and where *ΔTw* = *Tw* – *Tw*_0_ = 0, where *Tw_0_* is the natural twist of DNA, equal to the number of base pairs divided by the helical turn (≈ 10.5 bp per turn for bare DNA), and *ΔWr* = *Wr*, i.e. the torsionally relaxed conformation has *Wr* = 0.

Intercalation changes the intrinsic twist of the helix and, therefore, shifts the linking number difference at which the molecule is torsionally relaxed by *N*_bound_ · *Δθ*/360° where *N*_bound_ is the number of dye molecules bound. For a given plasmid, the linking number difference relative to the torsionally relaxed state is, therefore, given by

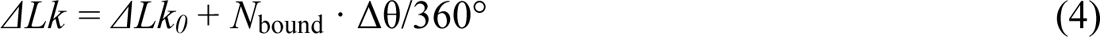

where *ΔLk_0_* is the linking number difference of the plasmid in the absence of intercalation. In general, excess linking number will partition into twist and writhe (Equation 3). For plasmids, it has been shown that the partitioning is independent of the magnitude (65) and sign (66) of the linking difference and is approximately 20% *Tw* and 80% *Wr*. We assume that this partioning between twist and writhe also holds in the presence of intercalators, such that

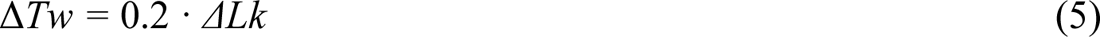

where *ΔLk* is given by Equation 4. In order to compute the torsional strain for a given initial linking difference *ΔLk_0_* and given number of intercalated molecules *N*_bound_, we convert the excess twist (Equation 5) to torque by taking into account the torsional stiffness of DNA:

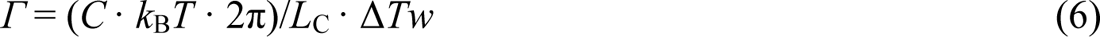

where *C* is the torsional stiffness of DNA in nm and *L*_C_ the contour length, which in turn depends on the number of molecules bound as *L*_C_ = *L*_C,0_ + *N*_bound_ ·*Δz*, where *L*_C,0_ is the contour length in the absence of intercalation, ≈ 0.34 nm per bp. The torsional stiffness of DNA has been measured using single-molecule methods (56,67,68), is independent of ionic strength (69), and reported values are in the range of *C* ≈ 100 nm (55–57). However, it is not well known whether or how *C* is altered by intercalation. Previous measurements using DNA in free solution using EtBr (27,70,71) have found lower values of the torsional stiffness in the range *C* ≈ 50 nm, which we take as a starting point for the EtBr data.

To determine the number of intercalated molecules per plasmid as a function of total ligand concentration *c*_total_ and DNA concentration *c*_DNA_ (typically expressed as the base pair concentration), we numerically solve the coupled Equations 1-6 using an iterative approach (Materials and Methods). The main output of the model is *N*_bound_. Assuming a linear relationship between the number of intercalated molecules and the fluorescence intensity, which we have previously found to hold for a large range of dye concentrations (15), the observed fluorescence intensity *I* is given by

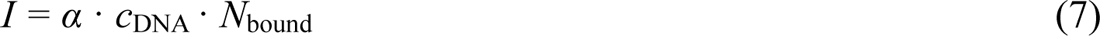

where *α* is a proportionality factor that depends on the quantum efficiency of the dye and the details of the experiments but is constant for a given intercalator and instrumental set up.

### DNA topology modulates intercalation

Our model for intercalation into topologically closed DNA predicts that binding can be increased or decreased by the global constraint, relatively to a torsionally relaxed (or linearized) plasmid. Starting with a negatively supercoiled plasmid, which is the form typically found *in vivo*, intercalation at low ligand concentration is increased relatively to a topologically open DNA, due to the negative torsional strain in the molecule (Figure 1F). As more and more molecules intercalate, the negative linking difference is compensated until the plasmid becomes torsionally relaxed, at which point binding to the topologically closed and open forms is the same. Finally, as the intercalator concentration is increased further, the closed plasmid becomes overwound, and the positive torsional strain hinders further intercalation. Therefore, at high intercalator concentration, our model predicts binding of fewer molecules to the closed compared to the open plasmid (Figure 1F).

To experimentally test the prediction of our model, we used plasmid DNA in both topologically constrained, negatively supercoiled form and in open circular and linear topologies (Figure 1A-C; Materials and Methods). We prepared mixtures with equal amounts of the three different DNA topologies to facilitate direct comparison on a gel. We then separated the mixtures on a gel (Figure 2A), imaged the gel, and quantified the band intensities to monitor the amount of intercalation (Supplementary Figure S3). While the open circle and linear topologies exhibit similar intensities, the topologically constrained species in comparison appears less bright on the gel for high EtBr concentrations (Supplementary Figure S4A). In contrast, for the lowest EtBr concentration, the supercoiled species exhibits a higher intensity than the other two species (Supplementary Figure S4B). We use our model to quantitatively account for the topology dependent intensities (Figure 2B,C). We find that for the experimental parameters used here, the number of intercalated molecules per plasmid is approximately independent of DNA concentration (Figure 2B). Consequently, the fluorescent intensity increases with DNA concentration (Figure 2C). At the lowest EtBr concentration (Figure 2C, black lines and symbols), more molecules bind to the supercoiled species (Figure 2C, lines and symbols with red highlighting) as compared to open circular and linear, while at the highest concentration (Figure 2C, light brown lines and symbols) intercalation is reduced for supercoiled compared to the other species. At the intermediate EtBr concentration (Figure 2C, brown lines and symbols) the topologically open and closed species bind EtBr similarly. The differences in binding between the different topologies are a consequence of the negative torsional strain at the lowest EtBr concentration and the positive strain for the highest concentration (Figure 2D). Averaging over the different DNA concentrations, we can quantitatively compare the relative enhancement or reduction of interaction for the topologically closed species compared to the open circular and linear species (Figure 2E) and find excellent agreement between our model and the experimental data.

**Figure 2.**
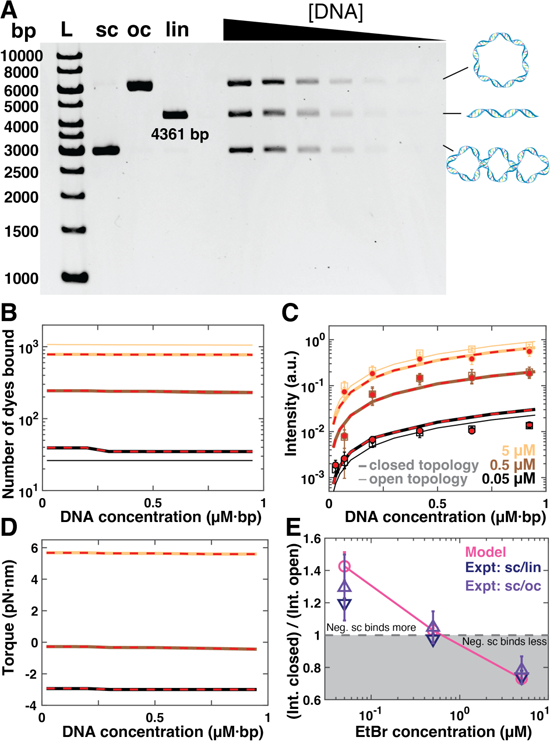
DNA topology dependent intercalation of ethidium bromide. **A)** Agarose gel stained with EtBr at a final concentration of 0.5 µM. Different DNA topologies are separated on the gel. L: DNA size ladders (1 kb gene ruler, Thermo Scientific, 5 μL). Lanes 2-4 are the stock solutions of negatively supercoiled, linear, and open circular DNA, respectively. Lanes 5-11 are equimolar mixtures of the three topologies, at different total DNA concentrations. **B)** Predicted number of intercalated molecules *N*_bound_ as function of DNA concentration for pBR322 DNA (4361 bp). Different colors correspond to different EtBr concentrations: from dark to light 0.05, 0.5, and 5 µM. Thin lines are for topologically open DNA (linear and open circular); thick lines with red highlights are for topologically closed DNA (supercoiled, here with supercoiling density σ = −5%, corresponding to *ΔLk*_0_ = −20 turns). The number of molecules bound is approximately independent of DNA concentration under the conditions investigated, but clearly dependents on EtBr concentration and DNA topology. **C)** Experimentally determined fluorescence intensity for supercoiled DNA (circles with red highlight) and linear DNA (squares) as a function EtBr and DNA concentration. Symbols are the mean and std from at least two gels. Lines are predictions of our binding model (same color code as in panel B), with the scale factor α as the only fitting parameter (Equation 7). **D)** Predicted torque in the plasmid from our model, same color code as in panel B. **E)** Relative fluorescence intensity of a topologically closed DNA relative to the topologically open constructs. Data points are obtained by averaging the different DNA concentration at the same EtBr condition. Triangles are experimental data from at least two independent gels. Magenta symbols are the prediction of or model.

### Topology dependent binding depends on initial topology and intercalator affinity

Having demonstrated that DNA topology can significantly alter DNA binding, we next use our quantitative model to explore how topology modulates intercalation. Starting with a negatively supercoiled plasmid (*ΔLk_0_*< 0) intercalation is increased at low intercalator concentrations and suppressed at high concentrations compared to an open topology (Figure 2E). A clear prediction of our model is that if the DNA is initially torsionally relaxed (or even positively supercoiled), intercalation should always be reduced for the closed topology compared to an open topology. We test this prediction experimentally by again preparing and separating DNA plasmids with different topologies, but now using a sample where the plasmid has been relaxed by topoisomerase treatment (Methods), such as that *ΔLk_0_* ≈ 0 (Figure 3A). As predicted, we find that EtBr intercalation is reduced for the closed topology (Figure 3C and Supplementary Figure S5), in excellent agreement with our model. More broadly, the relative effect of DNA topology depends both in the initial linking number *ΔLk_0_* and on the intercalator concentration (Figure 3E).

**Figure 3.**
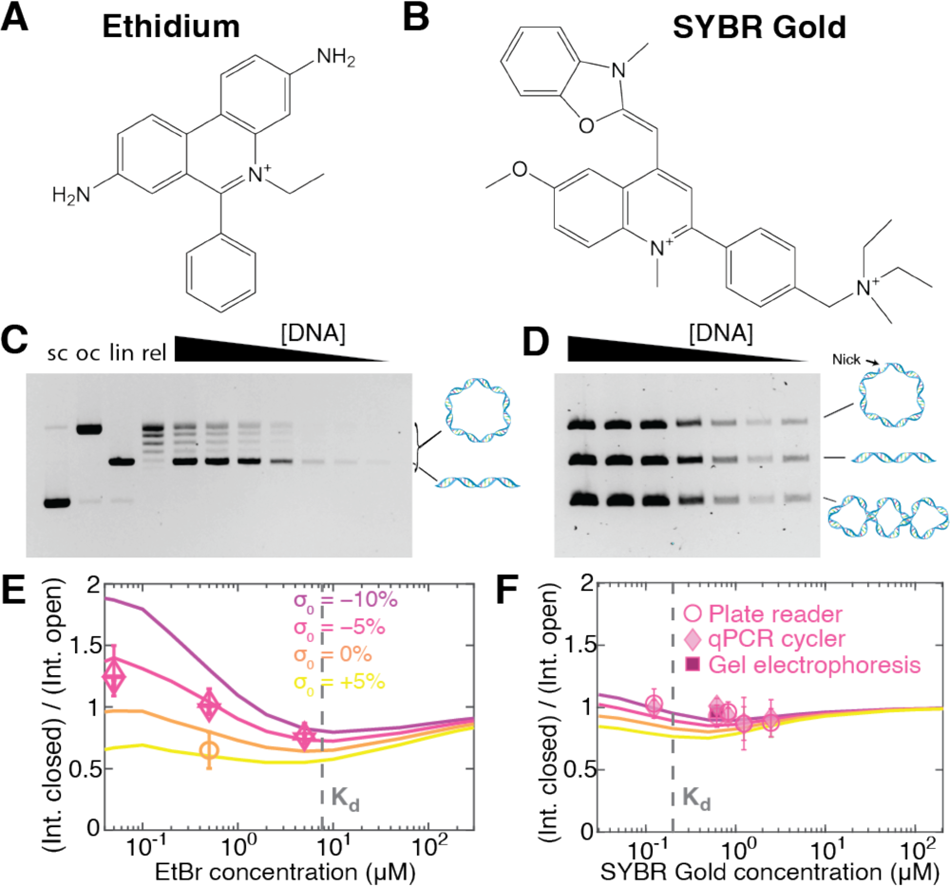
Topology dependent binding depends on concentrations and initial supercoiling density. **A)** Molecular structure of the intercalator ethidium bromide; taken from Ref. (81). **B)** Molecular structure of the intercalator SYBR Gold; taken from Ref. (15). **C)** Agarose gel with linear (“lin”) and topologically closed but initially torsionally relaxed DNA (“rel”), stained with EtBr at a final concentration of 0.5 µM. The linear DNA travels as a single band, while the relaxed DNA exhibits a topoisomer distribution. Lanes 5-11 are equimolar mixtures of the linear and relaxed DNA. **D)** Agarose gel with equimolar mixtures of open-circular, linear, and supercoiled DNA, stained with SYBR Gold at a final concentration of 0.6 µM. **E)** Relative binding to topologically closed DNA with different levels of initial supercoiling compared to topologically open constructs as a function of EtBr concentration. Colored lines are the predictions of our model. The vertical grey line indicates the *K*_d_ of EtBr. Symbols are the data from Figure 2E; further analysis is shown in Supplementary Figure S5. **F)** Same as in panel E for SYBR Gold. The data shown are from three different experimental modalities: fluorescent readout using a 96-well platereader, a qPCR cycler, and gel electrophoresis. Due to the much lower *K*_d_ for SYBR Gold compared to EtBr, the binding is almost independent of topology for the SYBR Gold conditions investigated.

Our model predicts that topology dependent binding is most pronounced at ligand concentration below the *K*_d_ (Figure 3E, the *K*_d_ value is indicated as a vertical line). At concentrations greater than the *K*_d_, binding saturates and the modulation by the torsional strain the in the topologically closed plasmid is predicted to only lead to small or negligible changes in binding compared to the torsionally relaxed forms. To test this prediction, we carried out measurements using the intercalator SYBR Gold (15,33,72), which has a much lower *K*_d_ (i.e. higher affinity) compared to EtBr (Table 1 and Figure 3B,D,F). Performing measurements with initially negatively supercoiled or relaxed plasmids at different SYBR Gold concentrations around and above its *K*_d_, we find that indeed the topologically closed and open constructs bind similar amounts of SYBR Gold, in quantitative agreement with our model (Figure 3F). To show the broad range of applications, we used three different assays to obtain fluorescence intensity data for SYBR Gold, namely gel electrophoresis, a well plate reader, and a qPCR cycler (see Methods for details).

Importantly, the almost topology-independent binding of DNA intercalators above their *K*_d_ is advantageous for assays that aim to quantitatively compare different DNA topologies, e.g., to monitor the products of integration or topoisomerization reactions (73–76). In particular, for SYBR Gold DNA staining is essentially unbiased by topology for dye concentrations in the range of 1-2 µM, which is the concentration range that we previously identified as optimal for achieving a linear relationship between the fluorescence signal and the amount of DNA present (15).

### Psoralen-based DNA crosslinking to quantify DNA supercoiling

Intercalators of the psoralen family can crosslink DNA upon irradiation with UV light (Figure 4A). They have been widely used in phototherapy (39) and to detect chromatin structure and the degree of DNA supercoiling *in vivo* (40–44). Often it is assumed that the amount of DNA crosslinking varies linearly with the supercoiling density σ (43,77). Our model for intercalation under the global constraint induced by topology accurately captures the relative binding of the psoralen compound TMP to supercoiled DNA vs. nicked DNA, determined from a radioactivity assay using ^3^H-labeled TMP (41) (Figure 4B). Similarly, our model correctly predicts the degree of crosslinking induced by TMP for different supercoiling densities (43) (Figure 4C and Supplementary Figure S6A). Importantly, the crosslinking conditions are chosen such that at most one crosslinking event per plasmid is induced, which means that only a small fraction of the intercalated TMP molecules reacts. For example, under the conditions of the data in Figure 4C, there are > 10 TMP molecules bound (Supplementary Figure S6B), but < 1 on average react. However, the fact that the crosslinking signal is well approximated by our binding model using a proportionality constant analogous to Equation 7, suggests that crosslinking is directly proportional to binding. While TMP binding is at least approximately linear with supercoiling density in the range investigated in Figure 4B and C, we note that a linear relationship is only an approximation to the intrinsic exponential dependence on torque and its validity is limited to relatively small supercoiling densities (Figure 4D).

**Figure 4.**
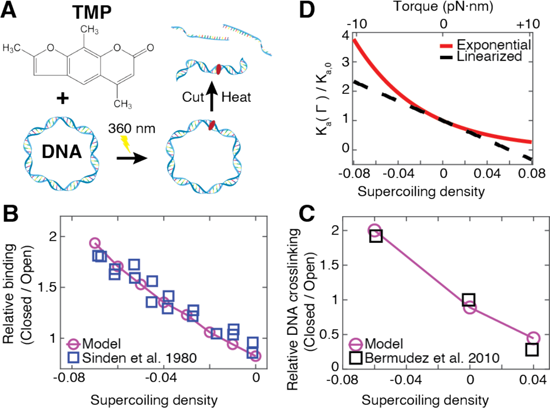
Topology-dependent DNA crosslinking by trimethylpsoralen. **A)** 4,5’,8-trimethylpsoralen (TMP) intercalates into DNA and causes DNA crosslinking upon UV irradiation. DNA supercoiling-dependent crosslinking is widely used to probe DNA supercoiling and chromatin conformations *in vivo*. **B)** Binding of TMP to supercoiled DNA plasmids with different initial supercoiling densities relative to open circular DNA. Experimental binding data are from Ref. (41) and were determined using the radioactivity of ^3^H-labeled TMP. **C)** Binding of TMP to supercoiled DNA plasmids. Experimental data are from Ref. (43) and were obtained by quantifying the amount of DNA crosslinking after irradiation. Experimental data are normalized to the data point at zero supercoiling density. The model in panels B and C uses the parameters in Table 1 and quantified binding to supercoiled relative to topologically open DNA. **D)** Dependence of the torque-dependent association constant (the inverse of the dissociation constant) on supercoiling density using Equations 2-6 (red solid line). The black dashed line shows the linearization of Equation 2, i.e., the approximation exp(−x) ≈ 1 – x, with x = *Γ·Δθ / k*_B_*T*.

### Single-molecule assay monitors topology dependent binding in real time

To explore consequences of topology dependent binding at the single-molecule level, we investigated ligand binding to DNA under a topological constraint via single-molecule fluorescence imaging. In our assay, we attached DNA via multiple biotin-streptavidin bonds at each end to a surface (Figure 5A), ensuring that the molecule is topologically constrained. Adding the intercalator SYTOX Orange enables us both to induce supercoiling in the DNA and to visualize the molecules using fluorescence imaging (35,38). To systematically study the effect of topology, we performed two different types of experiments (Materials and Methods). In the first case, we prepared negatively supercoiled DNA by first staining with the intercalative dye SYTOX Orange at a high concentration (250 nM), then attaching the DNA to the surface to topologically constrain it, and subsequently imaging at a lower concentration (50 nM). Following the reduction of SYTOX Orange concentration, plectonemic supercoils are clearly visible as bright spots that diffuse along the length of the DNA molecules (Figure 5B,C and Supplementary Movie S1).

**Figure 5.**
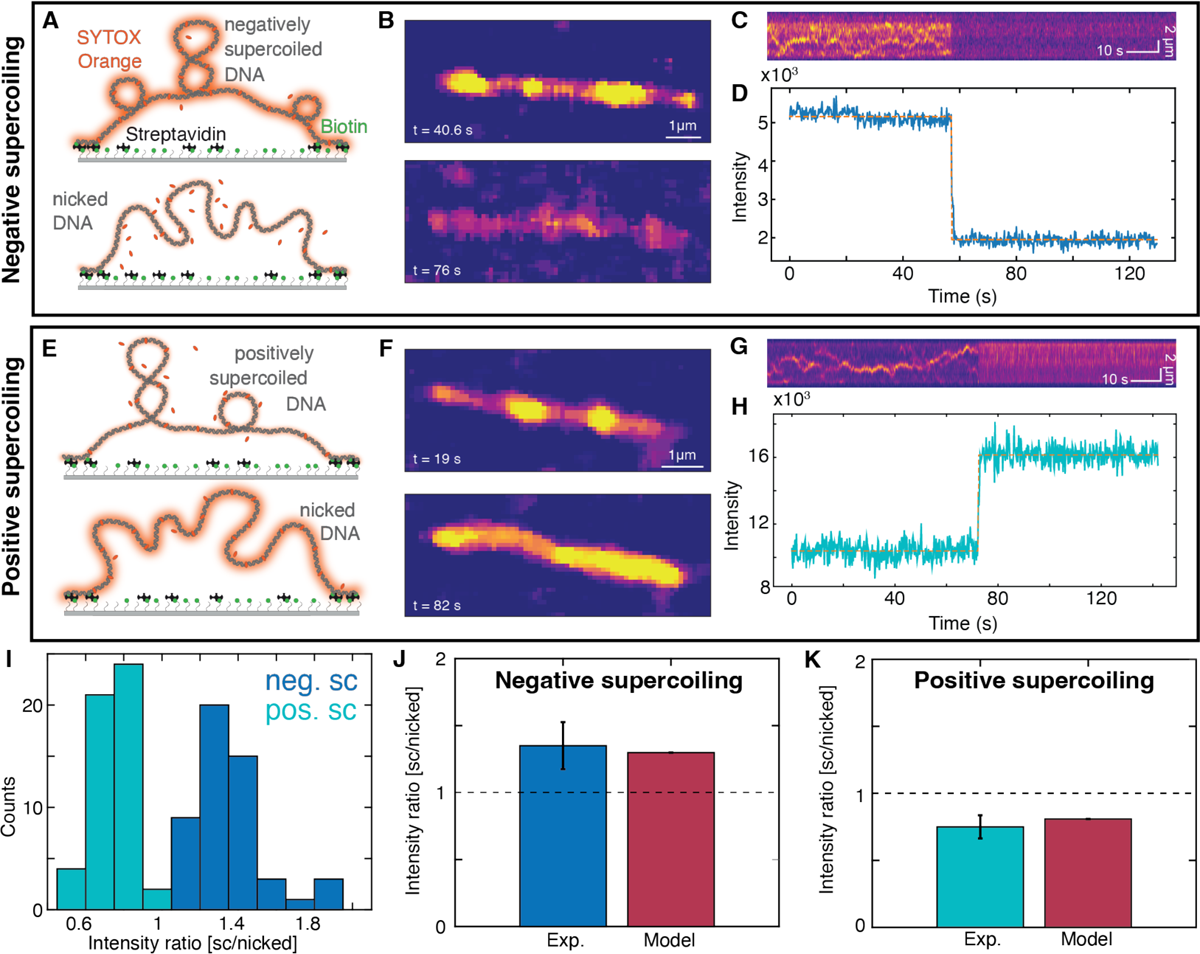
Single-molecule fluorescence assay to quantify topology dependent binding. A)-D): Negatively supercoiled DNA. A) Schematic representation of the experimental set up of the single-molecule fluorescence experiments. The SYTOX Orange-stained DNA is tethered at both its ends via multible biotin-streptavidin bonds to the surface. Top: negatively supercoiled DNA; bottom: nicked DNA. **B)** Fluorescence image snapshots at 40.6 s when the DNA is still negatively supercoiled and at 76 s after the DNA was nicked. **C)** Kymograph of SYTOX-Orange stained-DNA. When the DNA is nicked, the intensity decreases abruptly. **D)** Integrated fluorescence intensity of the kymograph shown in panel C. The dotted orange line is a fit of a two-state hidden Markov model to the data (92). **E)-H): Positively supercoiled DNA. E)** Schematic representation of the experimental lineup as in panel A only now the DNA is positively supercoiled before the nicking. **F)** Fluorescence image snapshots at 19 s when the DNA is still positively supercoiled and at 82 s after the DNA was nicked. **G)** Kymograph of SYTOX-Orange stained-DNA. When the DNA is nicked, the intensity increases abruptly. **H)** Fluorescence intensity of the kymograph shown in panel G. The dotted orange line is a fit (same as panel D) to the data. **I)** Intensity ratios before and after nicking for originally negatively supercoiled DNA (blue) and for for originally positively supercoiled DNA (turquoise). Averaging gives 1.35 ± 0.18 (N= 51, mean ± std; ratio sc/nicked) for originally negatively supercoiled DNA and 0.75 ± 0.09 (N= 51, mean ± std; ratio supercoiled/nicked) for originally positively supercoiled DNA. **J)** Comparison of the experimental intensity ratio to the value from theoretical modelling for for originally negatively supercoiled DNA. N= 51, **K)** Comparison of the experimental intensity ratio to the value from theoretical modelling for for originally positively supercoiled DNA. N= 51

During continuous observation and laser exposure, DNA molecule will nick, likely due to radicals generated by photochemical processes (78,79), which is usually an undesirable feature. However, here we use nicking upon illumination to our advantage since it enables us to observe both the supercoiled (closed) and later the nicked (open) topology of the same DNA molecule. Upon nicking, the fluorescence intensity suddenly decreases significantly (Figure 5D) and the bright spots indicative of plectonemic supercoils disappear (Figure 5B,C and Supplementary Movie S1). This is in line with our previous observations: The negative supercoiling helps intercalation, consequently, once the molecule nicks, less SYTOX Orange binds and the fluorescence intensity decreases.

For the second type of experiment, we prepared positively supercoiled DNA by attaching the DNA to the surface in the presence of a low SYTOX Orange concentration (25 nM; Figure 5E). We then increased the dye concentration (to 250 nM), but since positive supercoiling hinders intercalative binding to the DNA, the fluorescence intensity stays relatively low. Again, plectonemic supercoils appear as bright spots that diffusive along the DNA molecule (Figure 5F,G and Supplementary Movie S2). After the positively supercoiled molecule is nicked, we observe that the fluorescence intensity increases (Figure 5H). Importantly, our assay enables us to quantify the change in fluorescence intensity upon changes in topology by integrating the intensity over the entire molecule, before and after nicking (Figure 5D and H). We find a decrease in fluorescence intensity upon nicking for initially negatively supercoiled DNA of 1.35 ± 0.18 (mean ± std; ratio supercoiled/nicked) and an increase in fluorescence intensity upon nicking for initially positively supercoiled DNA of 0.75 ± 0.09 (mean ± std; ratio supercoiled/nicked) (Figure 5I).

To quantitatively model the changes in fluorescence upon nicking observed *in situ*, we used our model with the parameters reported by Biebricher *et al.* (34) for the binding site size *n*, elongation per dye *Δz*, and binding constant *K* taken in 100 mM NaCl, which approximately corresponds to the ionic strength in our experiments (40 mM Tris-HCl, 2.5 mM MgCl_2_, 65 mM KCl). The unwinding angle per intercalation event *Δθ* is not known for SYTOX Orange; we assume *Δθ* = 19.1°, which is the value for SYBR Gold (15) since the dyes are relatively similar dyes and also generally values in the range of about 20° are typical (15,80,81). Importantly, the model is applied here in two stages: We first compute the supercoiling density, relatively to relaxed, bare DNA and in the absence of intercalator, induced by attaching the DNA in the presence of 25 and 250 nM SYTOX Orange, which are σ = 2.7% and σ = 9.7%, respectively. We then compute binding to DNA and re-adjustment of the supercoiling level at the new SYTOX Orange concentrations used for imaging (250 and 50 nM, for which we find σ = +5.2% and σ = –3.9%), using the levels of supercoiling obtained in the first step as an input. For comparison, we compute binding to topologically open DNA, which enables us to calculate the changes in fluorescence upon nicking (Figure 5J,K). Here, we observe an excellent agreement between the predictions of our computed model and the experimentally observed changes in fluorescence intensity upon torsional relaxation in our single molecule experiments (Figure 5J,K).

### High-speed fluorescence tracking reveals binding dynamics

To quantitatively study the dynamics of intercalation into DNA under topological constraint, we performed single-molecule fluorescence imaging at a 20 ms frame rate (Figure 6), ten times faster than the data shown in Figure 5. By fitting a simple kinetic model to the fluorescence intensity traces (Figure 6A,B and E,F) at the transition between supercoiled and nicked DNA, we are able to determine the on- and off-rate of SYTOX Orange.

**Figure 6.**
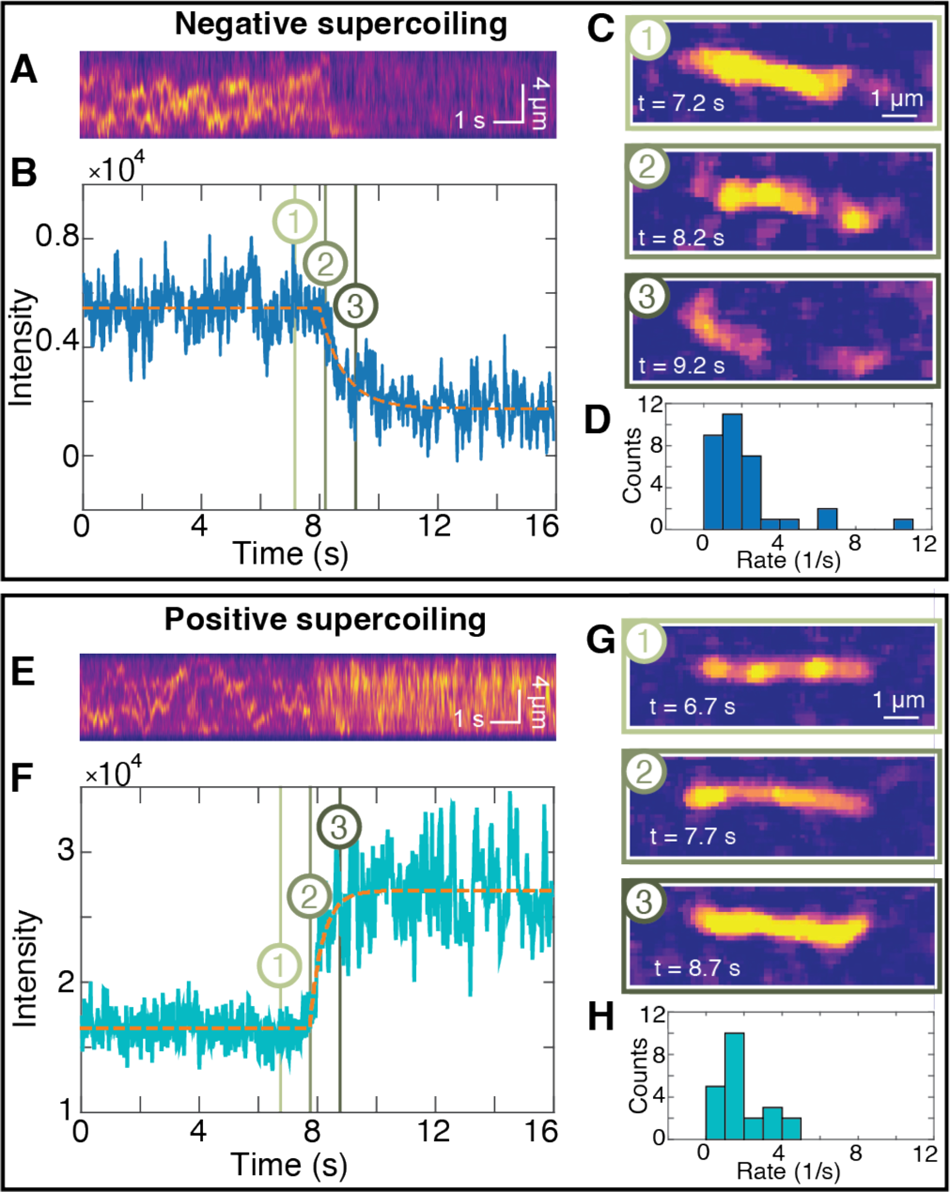
Probing the interplay of dye and supercoil dynamics. A)-D): Negatively supercoiled DNA. A) Kymograph of SYTOX-Orange stained-DNA. When the DNA is nicked, the intensity decreases abruptly. **B)** Integrated fluorescence intensity of the kymograph shown in panel A. The dotted orange line is a fit of the model shown in Equation 8 to the data. **C)** Fluorescence image snapshots at 7.2 s when the DNA is still negatively supercoiled, at 8.2 s when the nick occurs, and at 9.2 s after the DNA was nicked. Time points are indicated by matching numbers in panel B. **D)** Experimentally determined rates *k* for fluorescence decrease from the fits of Equation 8 to time traces from N = 32 independent measurements. The mean ± sem are (2.16 ± 0.38) 1/s. **E)-H): Positively supercoiled DNA. E)** Kymograph of SYTOX-Orange stained-DNA. When the DNA is nicked, the intensity increases abruptly. **F)** Integrated fluorescence intensity of the kymograph shown in panel E. The dotted orange line is a fit of Equation 8 to the data. **G)** Fluorescence image snapshots at 6.7 s when the DNA is still positively supercoiled, at 7.7 s when the nick occurs, and at 8.7 s after the DNA was nicked. Time points are indicated by matching numbers in panel F. **H)** Experimentally determined rates *k* for the increase in fluorescence after nicking from N = 22 independent measurements. The mean ± sem are (1.92 ± 0.26) 1/s.

Our kinetic model for the total fluorescence intensity of the initially supercoiled molecules, reads as follows:

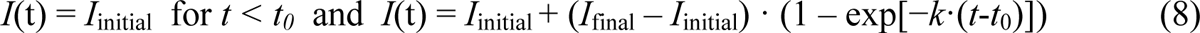

Where the initial intensity *I*_initial_, the final intensity *I*_final_, the rate *k*, and the time at which the intensity begins to change *t*_0_ are fitting parameters. For the DNA molecules that are negatively supercoiled prior to nicking, we find a reduction in fluorescence intensity with overall rate *k* = (2.16 ± 0.38) s^-1^ (mean ± sem from 32 traces; Figure 6D), which is close to the off-rate extrapolated to zero force reported by Biebricher et al. of (3.2 ± 0.8) s^-1^ using a single-molecule stretching assay (34). Conversely, starting with positively supercoiled DNA prior to nicking, we find an increase in intensity with an overall rate *k* = (1.92 ± 0.26) s^-1^ (mean ± sem from 22 traces; Figure 6H). Assuming that this increase is due to binding with a simple bimolecular association rate, we find an on-rate of (7.68 ± 1.3)·10^6^ M^-1^s^-1^ (since the SYTOX Orange concentration is constant at 250 nM in this case), again in excellent agreement to the on-rate reported by Biebricher *et al.* of (7.9 ± 2.6)·10^6^ M^-1^s^-1^. The very good agreement of our fitted overall rates with previously published on- and off-rates for SYTOX Orange suggests that binding and dissociation of the dyes and not the relaxation of torsional strain is rate limiting for the observed changes in fluorescence intensity. This is consistent with estimates from simulations that suggest that relaxation of torsional strain in DNA occurs on ∼µs time scales for ∼kbp DNA segments (82). The disappearance of the fluorescent spots (Figure 6C and G) allows us to estimate the dynamics of the writhe relaxation upon nicking, i.e., the time scale over which the plectonemic supercoils are resolved. We observe the disappearance of fluorescent spots over at least 3-4 frames (Supplementary Figure S7), suggesting that writhe relaxation occurs over ≥ 80 ms in our assay. The observed time scale for plectoneme disappearance is very similar to the lifetime of plectonemes before “hopping” events that have previously been detected by fluorescence imaging (83), suggesting that writhe relaxation occurs via “hopping” relaxation to a nick. Conversely, our measurements suggest that writhe relaxation does not occur predominantly by diffusion of plectonemes along the DNA to the nicking site, which would take on the order of τ ≈ *L^2^/D ≥* 3 s, where *L* is the length over which diffusion occurs (*L* ≥ 1 µm) and *D* the diffusion coefficient, which is ≤ 0.3 µm^2^/s (83).

## DISCUSSION

We have developed a quantitative model to describe DNA binding under topological constraints. Intriguingly, a global topological constraint can increase or decrease binding, depending on the concentration and binding regimes. This is an important difference to applied stretching forces, which similarly can modulate binding in an Arrhenius-like exponential dependence (21,84), but will bias binding in only one direction.

Using our model, we are able to elucidate the complex interplay between topology and ligand affinity. Using the well-characterized and widely used intercalators, SYBR Gold and ethidium bromide, we could show that topology-dependent binding depends on initial topology and intercalator affinity. In particular, we demonstrated that binding of DNA intercalators above their *K*_d_ is almost topology independent. With this, we can provide recommendations for optimal use of intercalative dyes to visualize DNA under a topological constraint: To avoid biases due to torque-dependent binding, a dye with a low *K*_d_ should be used, ideally at concentrations well above the *K*_d_. Specifically, SYBR Gold at a concentration of 1-2 µM satisfies this criterion. Importantly, SYBR Gold at 1-2 µM concentration is also the optimal concentration range to obtain a high signal as well as a linear relation between DNA amount and fluorescence intensity (15). In general, the choice of intercalator type and concentration range is crucial for reliably topology-unbiased DNA staining and quantification.

Moreover, our model can quantitatively account for observations made with the widely-used intercalator psoralen for UV-induced DNA crosslinking. Importantly, in applications where intercalation is used to detect supercoiling, a dye with a high *K*_d_ (i.e. low affinity) and large unwinding angle is desirable. Finally, we extend our model to quantitatively account for observations in single-molecule fluorescence measurements. This allows us to draw direct conclusions about the binding and dynamics of the DNA supercoiling.

Taken together, our work shows how combining theoretical modeling and multiple complementary experimental techniques can provide a highly quantitative and comprehensive view of DNA-ligand interactions under a global topological constraint. We anticipate our approach to be broadly applicable to other DNA binding agents and allow for reliable and unbiased detection and quantification of different topological states of DNA. In addition, the interplay of DNA binding and topology has important implications for the processing and regulation of genetic information *in vivo*. In the cell, DNA forms topological domains and e.g. advancing polymerases introduce torsional strains of different handedness (1,4,85–88). Together, these factors are expected to modulate binding to DNA, e.g. by nucleosomes and transcription factors, also in a cellular context. Our results provide a baseline to quantitatively investigate these complex processes in the future.

## DATA AVAILABILITY

All data are included in the article. Custom software that implements the topology dependent binding model written in Matlab is available freely in the repository YODA at https://doi.org/10.24416/UU01-YDNOBO.

## ACKNOWLEDGEMENTS

We thank Thomas Nicolaus for laboratory assistance, Felix Brandner for initial measurements, Patrick Kudella and Dieter Braun for help with qPCR cycler measurements, and Thorben Cordes for helpful discussions.

## REFERENCES

1. Liu, L.F. and Wang, J.C. (1987) Supercoiling of the DNA template during transcription. Proc Natl Acad Sci U S A, 84, 7024–7027.

2. Ubbink, J. and Odijk, T. (1999) Electrostatic-undulatory theory of plectonemically supercoiled DNA. Biophys J, 76, 2502–2519.

3. Vologodskii, A. (1994) DNA Extension Under the Action of an External Force. Macromolecules, 27, 5623–5625.

4. Koster, D.A., Crut, A., Shuman, S., Bjornsti, M.A. and Dekker, N.H. (2010) Cellular strategies for regulating DNA supercoiling: a single-molecule perspective. Cell, 142, 519–530.

5. Kouzine, F., Sanford, S., Elisha-Feil, Z. and Levens, D. (2008) The functional response of upstream DNA to dynamic supercoiling in vivo. Nat Struct Mol Biol, 15, 146–154.

6. Kouzine, F., Liu, J., Sanford, S., Chung, H.J. and Levens, D. (2004) The dynamic response of upstream DNA to transcription-generated torsional stress. Nat Struct Mol Biol, 11, 1092–1100.

7. Gilbert, N. and Allan, J. (2014) Supercoiling in DNA and chromatin. Current Opinion in Genetics & Development, 25, 15–21.

8. Corless, S. and Gilbert, N. (2016) Effects of DNA supercoiling on chromatin architecture. Biophysical Reviews, 8, 245–258.

9. Gebhardt, C., Lehmann, M., Reif, M.M., Zacharias, M. and Cordes, T. (2020) Molecular and spectroscopic characterization of green and red cyanine fluorophores from the Alexa Fluor and AF series. bioRxiv, 2020.2011.2013.381152.

10. Lerman, L.S. (1961) Structural considerations in the interaction of DNA and acridines. J Mol Biol, 3, 18–30.

11. Wang, J.G. (1974) The degree of unwinding of the DNA helix by ethidium: I. Titration of twisted PM2 DNA molecules in alkaline cesium chloride density gradients. Journal of Molecular Biology, 89, 783–801.

12. Long, E.C. and Barton, J.K. (1990) On demonstrating DNA intercalation. Accounts of Chemical Research, 23, 271–273.

13. Baguley, B. (1991) DNA intercalating anti-tumour agents. Anti-cancer drug design, 6, 1–35.

14. Langner, K.M., Kedzierski, P., Sokalski, W.A. and Leszczynski, J. (2006) Physical Nature of Ethidium and Proflavine Interactions with Nucleic Acid Bases in the Intercalation Plane. The Journal of Physical Chemistry B, 110, 9720–9727.

15. Kolbeck, P.J., Vanderlinden, W., Gemmecker, G., Gebhardt, C., Lehmann, M., Lak, A., Nicolaus, T., Cordes, T. and Lipfert, J. (2021) Molecular structure, DNA binding mode, photophysical properties and recommendations for use of SYBR Gold. Nucleic Acids Res, 49, 5143–5158.

16. Olson, W.K., Gorin, A.A., Lu, X.J., Hock, L.M. and Zhurkin, V.B. (1998) DNA sequence-dependent deformability deduced from protein-DNA crystal complexes. Proc Natl Acad Sci U S A, 95, 11163–11168.

17. Chou, F.C., Lipfert, J. and Das, R. (2014) Blind predictions of DNA and RNA tweezers experiments with force and torque. PLoS Comput Biol, 10, e1003756.

18. Skoruppa, E., Nomidis, S.K., Marko, J.F. and Carlon, E. (2018) Bend-Induced Twist Waves and the Structure of Nucleosomal DNA. Physical Review Letters, 121, 088101.

19. Vanderlinden, W., Lipfert, J., Demeulemeester, J., Debyser, Z. and De Feyter, S. (2014) Structure, mechanics, and binding mode heterogeneity of LEDGF/p75–DNA nucleoprotein complexes revealed by scanning force microscopy. Nanoscale, 6, 4611–4619.

20. Seol, Y. and Neuman, K.C. (2016) The Dynamic Interplay Between DNA Topoisomerases and DNA Topology. Biophys Rev, 8, 221–231.

21. Bell, G.I. (1978) Models for the specific adhesion of cells to cells. Science, 200, 618–627.

22. Evans, E. and Ritchie, K. (1997) Dynamic strength of molecular adhesion bonds. Biophys J, 72, 1541–1555.

23. Vladescu, I.D., McCauley, M.J., Nunez, M.E., Rouzina, I. and Williams, M.C. (2007) Quantifying force-dependent and zero-force DNA intercalation by single-molecule stretching. Nat Methods, 4, 517–522.

24. Lipfert, J., Klijnhout, S. and Dekker, N.H. (2010) Torsional sensing of small-molecule binding using magnetic tweezers. Nucleic Acids Res, 38, 7122–7132.

25. Salerno, D., Brogioli, D., Cassina, V., Turchi, D., Beretta, G.L., Seruggia, D., Ziano, R., Zunino, F. and Mantegazza, F. (2010) Magnetic tweezers measurements of the nanomechanical properties of DNA in the presence of drugs. Nucleic Acids Res, 38, 7089–7099.

26. Wang, J.C. (1974) The degree of unwinding of the DNA helix by ethidium. I. Titration of twisted PM2 DNA molecules in alkaline cesium chloride density gradients. J Mol Biol, 89, 783–801.

27. Selvin, P.R., Cook, D.N., Pon, N.G., Bauer, W.R., Klein, M.P. and Hearst, J.E. (1992) Torsional rigidity of positively and negatively supercoiled DNA. Science, 255, 82–85.

28. Sischka, A., Toensing, K., Eckel, R., Wilking, S.D., Sewald, N., Ros, R. and Anselmetti, D. (2005) Molecular mechanisms and kinetics between DNA and DNA binding ligands. Biophys J, 88, 404–411.

29. Hayashi, M. and Harada, Y. (2007) Direct observation of the reversible unwinding of a single DNA molecule caused by the intercalation of ethidium bromide. Nucleic Acids Res, 35, e125.

30. Celedon, A., Wirtz, D. and Sun, S. (2010) Torsional mechanics of DNA are regulated by small-molecule intercalation. J Phys Chem B, 114, 16929–16935.

31. Dikic, J. and Seidel, R. (2019) Anticooperative Binding Governs the Mechanics of Ethidium-Complexed DNA. Biophys J, 116, 1394–1405.

32. Tuma, R.S., Beaudet, M.P., Jin, X., Jones, L.J., Cheung, C.Y., Yue, S. and Singer, V.L. (1999) Characterization of SYBR Gold nucleic acid gel stain: a dye optimized for use with 300-nm ultraviolet transilluminators. Anal Biochem, 268, 278–288.

33. Cosa, G., Focsaneanu, K.S., McLean, J.R.N., McNamee, J.P. and Scaiano, J.C. (2001) Photophysical Properties of Fluorescent DNA-dyes Bound to Single- and Double-stranded DNA in Aqueous Buffered Solution¶. Photochemistry and Photobiology, 73, 585–599.

34. Biebricher, A.S., Heller, I., Roijmans, R.F., Hoekstra, T.P., Peterman, E.J. and Wuite, G.J. (2015) The impact of DNA intercalators on DNA and DNA-processing enzymes elucidated through force-dependent binding kinetics. Nat Commun, 6, 7304.

35. Ganji, M., Kim, S.H., van der Torre, J., Abbondanzieri, E. and Dekker, C. (2016) Intercalation-Based Single-Molecule Fluorescence Assay To Study DNA Supercoil Dynamics. Nano Letters, 16, 4699–4707.

36. King, G.A., Biebricher, A.S., Heller, I., Peterman, E.J.G. and Wuite, G.J.L. (2018) Quantifying Local Molecular Tension Using Intercalated DNA Fluorescence. Nano Letters, 18, 2274–2281.

37. Ganji, M., Shaltiel, I.A., Bisht, S., Kim, E., Kalichava, A., Haering, C.H. and Dekker, C. (2018) Real-time imaging of DNA loop extrusion by condensin. Science, 360, 102–105.

38. Kim, S.H., Ganji, M., Kim, E., van der Torre, J., Abbondanzieri, E. and Dekker, C. (2018) DNA sequence encodes the position of DNA supercoils. eLife, 7, e36557.

39. Lapolla, W., Yentzer, B.A., Bagel, J., Halvorson, C.R. and Feldman, S.R. (2011) A review of phototherapy protocols for psoriasis treatment. Journal of the American Academy of Dermatology, 64, 936–949.

40. Cech, T. and Pardue, M.L. (1977) Cross-linking of DNA with trimethylpsoralen is a probe for chromatin structure. Cell, 11, 631–640.

41. Sinden, R.R., Carlson, J.O. and Pettijohn, D.E. (1980) Torsional tension in the DNA double helix measured with trimethylpsoralen in living E. coli cells: Analogous measurements in insect and human cells. Cell, 21, 773–783.

42. Cimino, G.D., Gamper, H.B., Isaacs, S.T. and Hearst, J.E. (1985) Psoralens as photoactive probes of nucleic acid structure and function: organic chemistry, photochemistry, and biochemistry. Annual Review of Biochemistry, 54, 1151–1193.

43. Bermúdez, I., García-Martínez, J., Pérez-Ortín, J.E. and Roca, J. (2010) A method for genome-wide analysis of DNA helical tension by means of psoralen–DNA photobinding. Nucleic Acids Res, 38, e182–e182.

44. Naughton, C., Avlonitis, N., Corless, S., Prendergast, J.G., Mati, I.K., Eijk, P.P., Cockroft, S.L., Bradley, M., Ylstra, B. and Gilbert, N. (2013) Transcription forms and remodels supercoiling domains unfolding large-scale chromatin structures. Nature Structural & Molecular Biology, 20, 387–395.

45. Yamashita, Y., Fujii, N., Murakata, C., Ashizawa, T., Okabe, M. and Nakano, H. (1992) Induction of mammalian DNA topoisomerase I mediated DNA cleavage by antitumor indolocarbazole derivatives. Biochemistry, 31, 12069–12075.

46. Webb, M.R. and Ebeler, S.E. (2003) A gel electrophoresis assay for the simultaneous determination of topoisomerase I inhibition and DNA intercalation. Analytical Biochemistry, 321, 22–30.

47. Maxwell, A., Burton, N.P. and O’Hagan, N. (2006) High-throughput assays for DNA gyrase and other topoisomerases. Nucleic Acids Res, 34, e104.

48. Kolbeck, P.J., Dass, M., Martynenko, I.V., van Dijk-Moes, R.J.A., Brouwer, K.J.H., van Blaaderen, A., Vanderlinden, W., Liedl, T. and Lipfert, J. (2023) DNA Origami Fiducial for Accurate 3D Atomic Force Microscopy Imaging. Nano Letters.

49. Konrad, S.F., Vanderlinden, W. and Lipfert, J. (2021) A High-throughput Pipeline to Determine DNA and Nucleosome Conformations by AFM Imaging. Bio-protocol, 11, e4180.

50. Bussiek, M., Mücke, N. and Langowski, J. (2003) Polylysine-coated mica can be used to observe systematic changes in the supercoiled DNA conformation by scanning force microscopy in solution. Nucleic Acids Res, 31, e137–e137.

51. Chandradoss, S.D., Haagsma, A.C., Lee, Y.K., Hwang, J.-H., Nam, J.-M. and Joo, C. (2014) Surface Passivation for Single-molecule Protein Studies. JoVE, e50549.

52. Tišma, M., Panoukidou, M., Antar, H., Soh, Y.-M., Barth, R., Pradhan, B., Barth, A., van der Torre, J., Michieletto, D., Gruber, S. et al. (2022) ParB proteins can bypass DNA-bound roadblocks via dimer-dimer recruitment. Science Advances, 8, eabn3299.

53. Aitken, C.E., Marshall, R.A. and Puglisi, J.D. (2008) An Oxygen Scavenging System for Improvement of Dye Stability in Single-Molecule Fluorescence Experiments. Biophysical Journal, 94, 1826–1835.

54. Pradhan, B., Kanno, T., Umeda Igarashi, M., Loke, M.S., Baaske, M.D., Wong, J.S.K., Jeppsson, K., Björkegren, C. and Kim, E. (2023) The Smc5/6 complex is a DNA loop-extruding motor. Nature, 616, 843–848.

55. Bryant, Z., Stone, M.D., Gore, J., Smith, S.B., Cozzarelli, N.R. and Bustamante, C. (2003) Structural transitions and elasticity from torque measurements on DNA. Nature, 424, 338–341.

56. Lipfert, J., Kerssemakers, J.W., Jager, T. and Dekker, N.H. (2010) Magnetic torque tweezers: measuring torsional stiffness in DNA and RecA-DNA filaments. Nat Methods, 7, 977–980.

57. Mosconi, F., Allemand, J.F., Bensimon, D. and Croquette, V. (2009) Measurement of the torque on a single stretched and twisted DNA using magnetic tweezers. Phys Rev Lett, 102, 078301.

58. Ali, M., Lipfert, J., Seifert, S., Herschlag, D. and Doniach, S. (2010) The ligand-free state of the TPP riboswitch: a partially folded RNA structure. J Mol Biol, 396, 153–165.

59. Golub, G.H. and Van Loan, C.F. (2013) Matrix computations. JHU press.

60. McGhee, J.D. and Hippel, P.H.v. (1974) Theoretical Aspects of DNA-Protein Interactions: Co-operative and Non-co-operative Binding of Large Ligands to a One-dimensional Homogeneous Lattice. J. Mol. Biol., 86, 469–489.

61. Wang, J.C. (1969) Variation of the average rotation angle of the DNA helix and the superhelical turns of covalently closed cyclic λ DNA. Journal of Molecular Biology, 43, 25–39.

62. Bauer, W.R. (1978) Structure and reactions of closed duplex DNA. Annual review of biophysics and bioengineering, 7, 287–313.

63. Anderson, P. and Bauer, W. (1978) Supercoiling in closed circular DNA: dependence upon ion type and concentration. Biochemistry, 17, 594–601.

64. White, J.H. (1969) Self-Linking and Gauss-Integral in higher dimensions. Am. J. Math., 91, 693-&.

65. Boles, T.C., White, J.H. and Cozzarelli, N.R. (1990) Structure of plectonemically supercoiled DNA. J Mol Biol, 213, 931–951.

66. Brouns, T., De Keersmaecker, H., Konrad, S.F., Kodera, N., Ando, T., Lipfert, J., De Feyter, S. and Vanderlinden, W. (2018) Free Energy Landscape and Dynamics of Supercoiled DNA by High-Speed Atomic Force Microscopy. ACS Nano, 12, 11907–11916.

67. Moroz, J.D. and Nelson, P. (1997) Torsional directed walks, entropic elasticity, and DNA twist stiffness. Proc Natl Acad Sci U S A, 94, 14418–14422.

68. Kriegel, F., Ermann, N. and Lipfert, J. (2017) Probing the mechanical properties, conformational changes, and interactions of nucleic acids with magnetic tweezers. J Struct Biol, 197, 26–36.

69. Kriegel, F., Ermann, N., Forbes, R., Dulin, D., Dekker, N.H. and Lipfert, J. (2017) Probing the salt dependence of the torsional stiffness of DNA by multiplexed magnetic torque tweezers. Nucleic Acids Res, 45, 5920–5929.

70. Fujimoto, B.S. and Schurr, J.M. (1990) Dependence of the torsional rigidity of DNA on base composition. Nature, 344, 175–177.

71. Heath, P.J., Clendenning, J.B., Fujimoto, B.S. and Schurr, J.M. (1996) Effect of bending strain on the torsion elastic constant of DNA. J Mol Biol, 260, 718–730.

72. Tuma, R.S., Beaudet, M.P., Jin, X., Jones, L.J., Cheung, C.-Y., Yue, S. and Singer, V.L. (1999) Characterization of SYBR Gold Nucleic Acid Gel Stain: A Dye Optimized for Use with 300-nm Ultraviolet Transilluminators. Analytical Biochemistry, 268, 278–288.

73. Vanderlinden, W., Brouns, T., Walker, P.U., Kolbeck, P.J., Milles, L.F., Ott, W., Nickels, P.C., Debyser, Z. and Lipfert, J. (2019) The free energy landscape of retroviral integration. Nature Communications, 10, 4738.

74. Maxwell, A., Burton, N.P. and O’Hagan, N. (2006) High-throughput assays for DNA gyrase and other topoisomerases. Nucleic Acids Res, 34, e104–e104.

75. Molloy, M.J., Hall, V.S., Bailey, S.I., Griffin, K.J., Faulkner, J. and Uden, M. (2004) Effective and robust plasmid topology analysis and the subsequent characterization of the plasmid isoforms thereby observed. Nucleic Acids Res, 32, e129–e129.

76. Wang, M., Liu, J.-K., Gao, T., Xu, L.-L., Zhang, X.-X., Nie, J.-H., Li, Y. and Chen, H.-X. (2022) A platform method for plasmid isoforms analysis by capillary gel electrophoresis. Electrophoresis, 43, 1174–1182.

77. (2016) In Bienz, S., Bigler, L. and Fox, T. (eds.), Spektroskopische Methoden in der organischen Chemie. 9. Auflage ed. Georg Thieme Verlag.

78. Tycon, M.A., Dial, C.F., Faison, K., Melvin, W. and Fecko, C.J. (2012) Quantification of dye-mediated photodamage during single-molecule DNA imaging. Analytical Biochemistry, 426, 13–21.

79. Paillous, N. and Vicendo, P. (1993) Mechanisms of photosensitized DNA cleavage. Journal of Photochemistry and Photobiology B: Biology, 20, 203–209.

80. Vanderlinden, W., Kolbeck, P.J., Frederickx, W., Konrad, S.F., Nicolaus, T., Lampe, C., Urban, A.S., Moucheron, C. and Lipfert, J. (2019) Ru(TAP)32+ uses multivalent binding to accelerate and constrain photo-adduct formation on DNA. Chemical Communications, 55, 8764–8767.

81. Lipfert, J., Klijnhout, S. and Dekker, N.H. (2010) Torsional sensing of small-molecule binding using magnetic tweezers. Nucleic Acids Res, 38, 7122–7132.

82. Fosado, Y.A.G., Michieletto, D., Brackley, C.A. and Marenduzzo, D. (2021) Nonequilibrium dynamics and action at a distance in transcriptionally driven DNA supercoiling. Proceedings of the National Academy of Sciences, 118, e1905215118.

83. van Loenhout, M.T., de Grunt, M.V. and Dekker, C. (2012) Dynamics of DNA supercoils. Science, 338, 94–97.

84. Vladescu, I.D., McCauley, M.J., Nuñez, M.E., Rouzina, I. and Williams, M.C. (2007) Quantifying force-dependent and zero-force DNA intercalation by single-molecule stretching. Nature Methods, 4, 517–522.

85. Finzi, L. and Dunlap, D. (2016) Supercoiling biases the formation of loops involved in gene regulation. Biophys Rev, 8, 65–74.

86. Lipfert, J., van Oene, M.M., Lee, M., Pedaci, F. and Dekker, N.H. (2015) Torque spectroscopy for the study of rotary motion in biological systems. Chem Rev, 115, 1449–1474.

87. Naumova, N., Imakaev, M., Fudenberg, G., Zhan, Y., Lajoie, B.R., Mirny, L.A. and Dekker, J. (2013) Organization of the mitotic chromosome. Science, 342, 948–953.

88. Postow, L., Hardy, C.D., Arsuaga, J. and Cozzarelli, N.R. (2004) Topological domain structure of the Escherichia coli chromosome. Genes Dev, 18, 1766–1779.

89. Diekmann, J., Theves, I., Thom, K.A. and Gilch, P. (2020) Tracing the Photoaddition of Pharmaceutical Psoralens to DNA. Molecules, 25, 5242.

90. Wiesehahn, G. and Hearst, J.E. (1978) DNA unwinding induced by photoaddition of psoralen derivatives and determination of dark-binding equilibrium constants by gel electrophoresis. Proc Natl Acad Sci U S A, 75, 2703–2707.

91. Lercher, L., McDonough, M.A., El-Sagheer, A.H., Thalhammer, A., Kriaucionis, S., Brown, T. and Schofield, C.J. (2014) Structural insights into how 5-hydroxymethylation influences transcription factor binding. Chemical Communications, 50, 1794–1796.

92. Liu, H. and Shima, T. (2021) A fast and objective hidden Markov modeling for accurate analysis of biophysical data with numerous states. bioRxiv, 2021.2005.2030.446337.

