## Supplementary Information for "Topology-dependent DNA binding"

| **Equimolar DNA mix (μL)** | **TAE buffer (μL)** | **Gel loading dye (μL)** | **[DNA] (μM · bp)** |
|---|---|---|---|
| 20 | / | / | 0.9231 |
| 15 | 5 | 1 | 0.6585 |
| 10 | 10 | 2 | 0.4185 |
| 5 | 15 | 3 | 0.2000 |
| 2 | 20 | 4 | 0.0708 |
| 1 | 25 | 5 | 0.0298 |
| 0.5 | 25 | 5 | 0.0151 |

**Table S1: DNA dilution series for gel electrophoresis assays.**

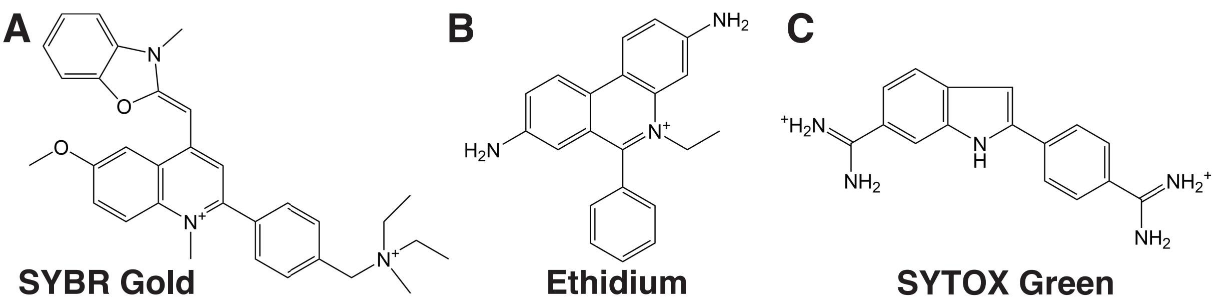


**Figure S1. Molecular structure of the three intercalators (A) SYBR Gold, (B) Ethidium, (C) SYTOX Green.** The SYBR Gold structure was taken from Reference (1), the Ethidium structure from reference (2). Since the structure of SYTOX Orange, the dye used for the single-molecule fluorescence assay in this work, is unknown, we show the structure of SYTOX Green (taken from reference (3)) here.

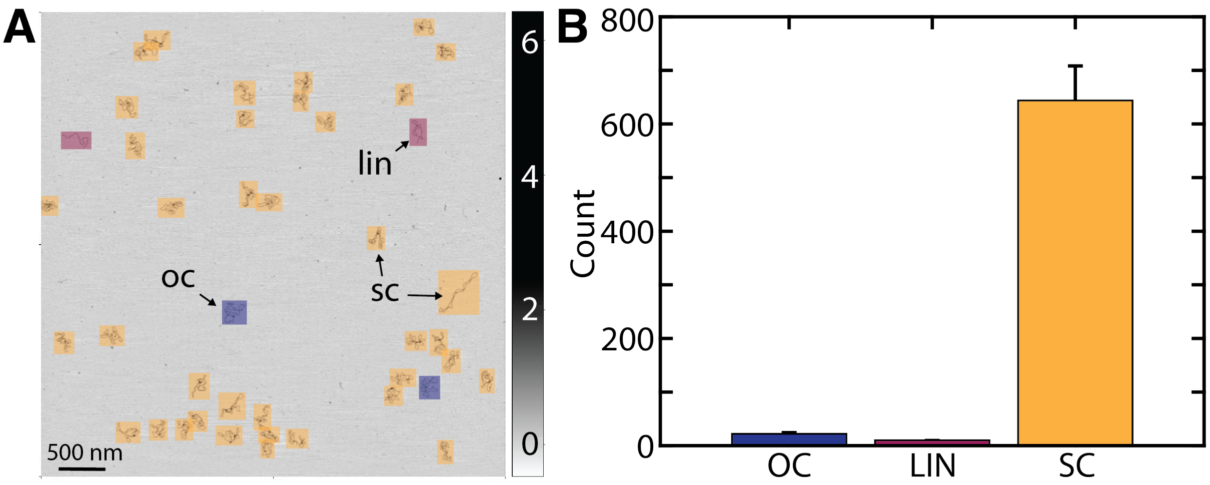


**Figure S2. DNA topology analysis and quality control via AFM imaging. A)** AFM height image of supercoiled pBR322 DNA at a concentration of 1 ng/μL deposited on PLL mica after drying in air. The different topologies are indicated with different colors, oc DNA in blue, lin DNA in red, and sc DNA in yellow. Z-ranges are indicated in nm by the scale bar on the right. **B)** Topology analysis from AFM experiments of pBR322 DNA that was also used as supercoiled DNA for gel experiments. From a total of 676 molecules, 95% are supercoiled, 3% open circular, and 2% linear. Error bars are from counting statistics.

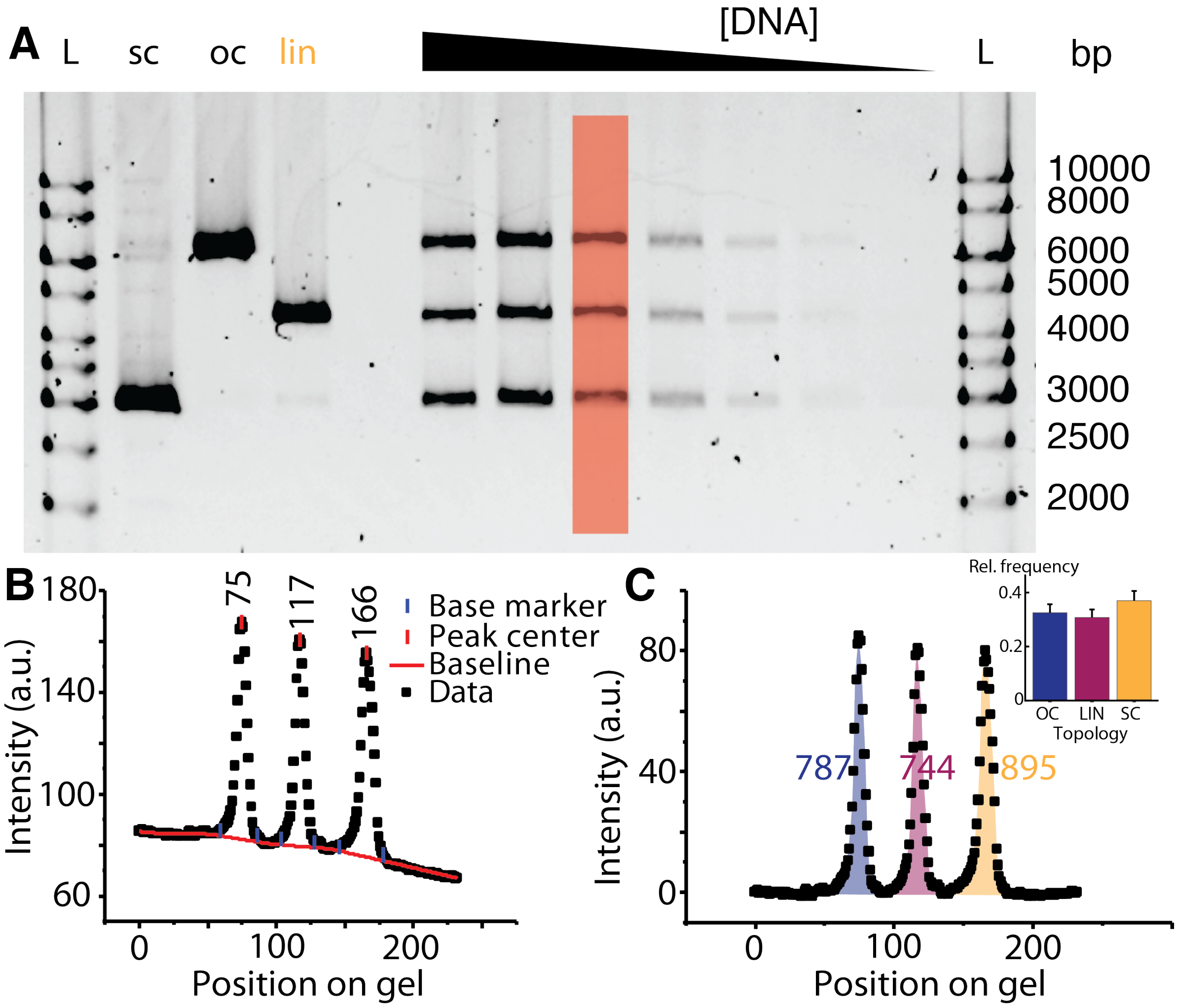


**Figure S3. Analysis of gel electrophoresis data. A)** Agarose gel stained with SYBR Gold at a final concentration of 0.315 µM. Different DNA topologies are separated on the gel. L: DNA size ladders (1 kb gene ruler, Thermo Scientific, 5 µL). Lanes 2-4 are the stock solutions of the supercoiled, linear, and open circular DNA, respectively. Lanes 6-12 are equimolar mixtures of the three topologies, at different total DNA concentrations. For quantitative analysis, individual lanes are selected (as highlighted in red) to create intensity profiles. **B)** Line profile of the area highlighted in red in panel A. Using the software Origin, a baseline is set and the peaks corresponding to the three topologies (oc, lin, sc) are detected automatically. **C)** Same data as in panel B, only baseline corrected. The area under the peaks (which corresponds to the fluorescence intensity of the individual topologies) is calculated and used to determine the intensity ratios of the three topological states. Inset: integrated and normalized intensities of the individual peaks shown in panel C.

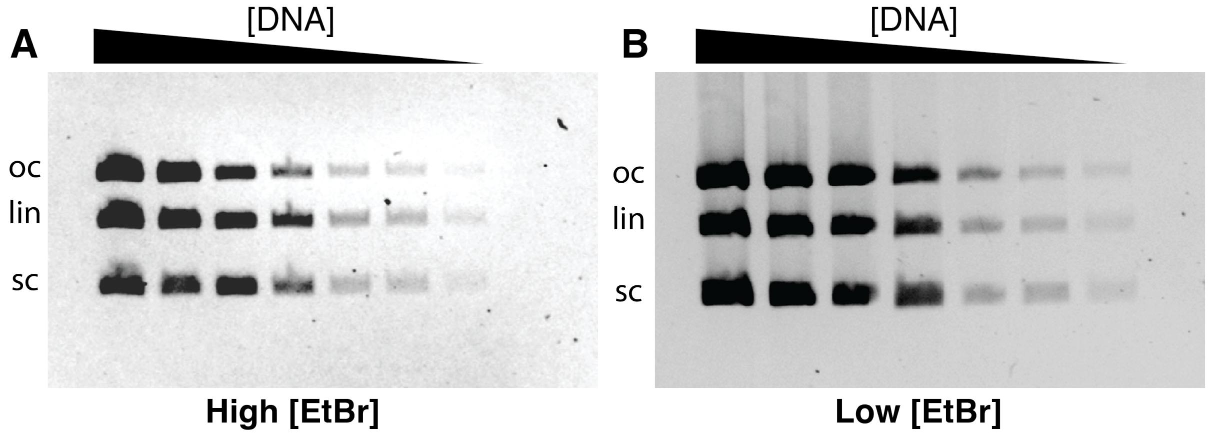


**Figure S4. Agarose gel stained with EtBr at a final concentration of 5 µM (A) and 0.05 µM (B), respectively.** Different DNA topologies are separated on the gel. Lanes 1-7 are equimolar mixtures of the three topologies, at different total DNA concentrations.

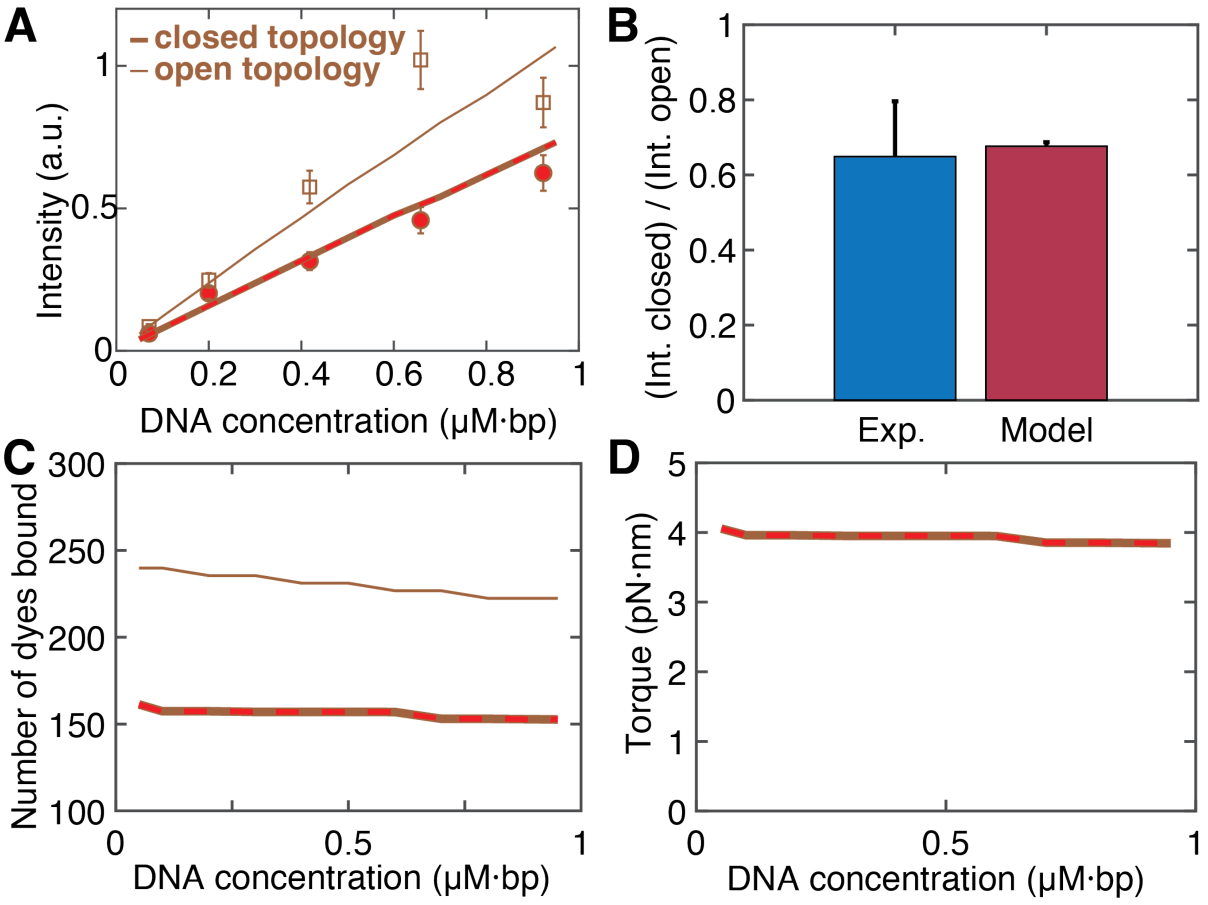


**Figure S5. Topology dependent binding of ethidium bromide. A)** Experimentally determined fluorescence intensity for topologically closed DNA (circles with red highlight) and lin DNA (squares) as a function of DNA concentration. Lines are predictions of our binding model (same color code as in Figure 2), with the scale factor α as the only fitting parameter (Equation 7). The EtBr concentration is 0.5 µM. **B)** Relative fluorescence intensity of a topologically closed DNA with $\Delta Lk_0$=0 relative to the topologically open DNA. The experimental data are the mean and std over different DNA concentration. **C)** Predicted number of intercalated molecules $N_{\mathrm{bound}}$ as function of DNA concentration for pBR322 DNA (4361 bp) with $\Delta Lk_0$=0. Same color code as Figure 2. The number of molecules bound depends only weakly on DNA concentration under the conditions investigated but is clearly reduced for the closed DNA topology compared to linear DNA. **D)** Predicted torque in the plasmid from our model, same color code as in panel A,C.

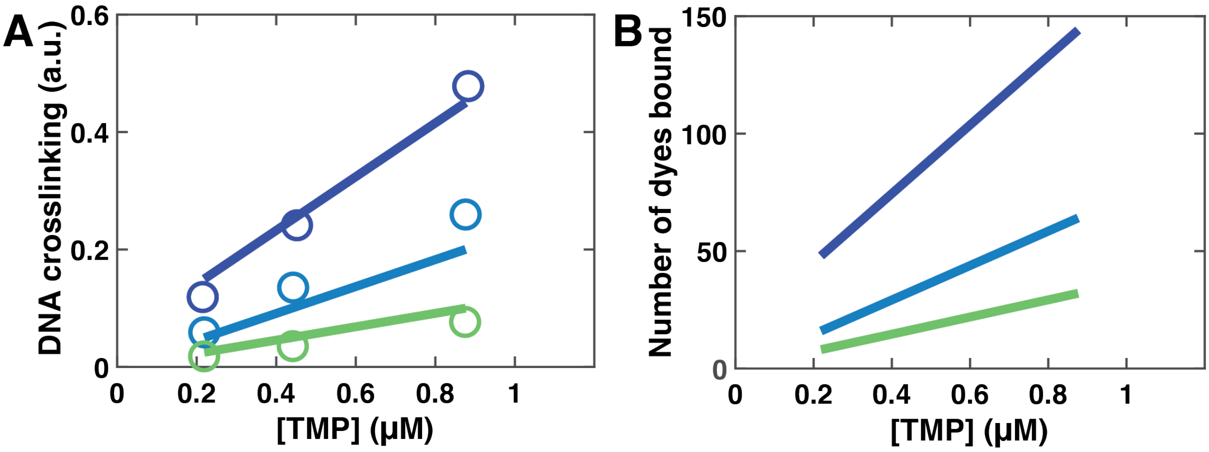


**Figure S6. Model prediction for TMP binding to plasmid DNA. A)** Number of DNA crosslinks as a function of TMP concentration for DNA plasmids with different initial supercoiling density (from blue to green, top to bottom: $\sigma = -0.06, 0, +0.04$). Circles are the experimental data from Ref. (4). Solid lines are the prediction of our model using the parameters in Table 1. B) Number of dyes bound as a function of TMP concentration. Same colour code as in panel A.

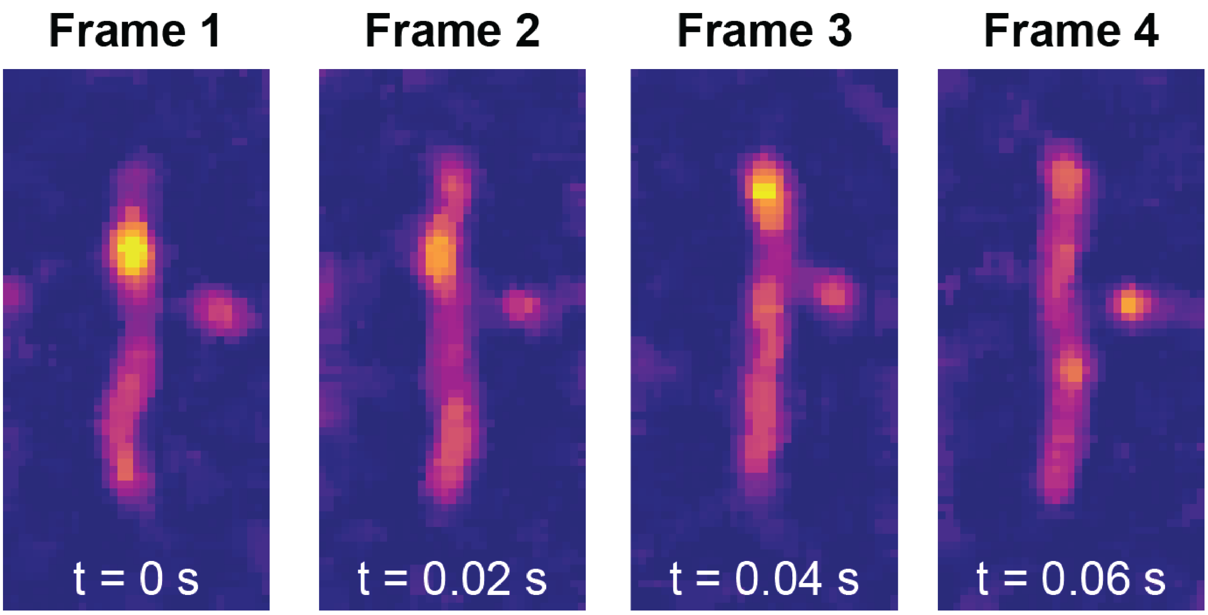


**Figure S7. Subsequent fluorescence image snapshots at the transition between negatively supercoiled and nicked DNA.** We observe that the fluorescent spots disappear over 4 frames (20 ms frame rate), suggesting that writhe relaxation occurs within approximately 80 ms.

**SUPPLEMENTARY REFERENCES**


1. Kolbeck, P.J., Vanderlinden, W., Gemmecker, G., Gebhardt, C., Lehmann, M., Lak, A., Nicolaus, T., Cordes, T. and Lipfert, J. (2021) Molecular structure, DNA binding mode, photophysical properties and recommendations for use of SYBR Gold. *Nucleic Acids Res*, **49**, 5143-5158.
2. Lipfert, J., Klijnhout, S. and Dekker, N.H. (2010) Torsional sensing of small-molecule binding using magnetic tweezers. *Nucleic Acids Res*, **38**, 7122-7132.
3. Wright, D.A. and Welschmeyer, N.A. (2015) Establishing benchmarks in compliance assessment for the ballast water management convention by port state control. *Journal of Marine Engineering & Technology*, **14**, 9-18.
4. Bermúdez, I., García-Martínez, J., Pérez-Ortín, J.E. and Roca, J. (2010) A method for genome-wide analysis of DNA helical tension by means of psoralen–DNA photobinding. *Nucleic Acids Res*, **38**, e182-e182.
